# ARHGEF26 enhances *Salmonella* invasion and inflammation in cells and mice

**DOI:** 10.1101/2020.12.30.424811

**Authors:** Jeffrey S. Bourgeois, Liuyang Wang, Monica I. Alvarez, Jeffrey Everitt, Sahezeel Awadia, Erika S. Wittchen, Rafael Garcia-Mata, Dennis C. Ko

**Affiliations:** Department of Molecular Genetics and Microbiology, School of Medicine, Duke University, Durham, NC, USA; University Program in Genetics and Genomics, Duke University, Durham, NC, USA; Department of Pathology, Duke University Medical Center, Durham, NC, USA; Department of Biological Sciences, University of Toledo, Toledo, OH, USA; Department of Cell Biology and Physiology, University of North Carolina, Chapel Hill, NC, USA; Division of Infectious Diseases, Department of Medicine, School of Medicine, Duke University, Durham, NC, USA

## Abstract

*Salmonella* hijack host machinery in order to invade cells and establish infection. While considerable work has described the role of host proteins in invasion, much less is known regarding how natural variation in these invasion-associated host proteins affects *Salmonella* pathogenesis. Here we leveraged a candidate cellular GWAS screen to identify natural genetic variation in the *ARHGEF26 (Rho Guanine Nucleotide Exchange Factor 26*) gene that renders lymphoblastoid cells susceptible to *Salmonella* Typhi and Typhimurium invasion. Experimental follow-up redefined ARHGEF26’s role in *Salmonella* epithelial cell invasion, identified serovar specific interactions, implicated ARHGEF26 in SopE-mediated invasion, and revealed that the ARHGEF26-associated proteins DLG1 and SCRIB facilitate *S*. Typhi uptake. Importantly, we show that ARHGEF26 plays a critical role in *S*. Typhimurium pathogenesis by contributing to bacterial burden in the enteric fever murine model, as well as inflammation in the gastroenteritis infection model. The impact of *ARHGEF26* on inflammation was also seen in cells, as knockdown reduced IL-8 production in HeLa cells. Together, these data reveal pleiotropic roles for ARHGEF26 function during infection and highlight that many of the interactions that occur during infection that are thought to be well understood likely have underappreciated complexity.

**Author Summary:** During infection, *Salmonella* manipulates host cells into engulfing the bacteria and establishing an intracellular niche. While many studies have identified genes involved in different stages of this *Salmonella* invasion process, few studies have examined how differences between human hosts contribute to infection susceptibility. Here we leveraged a candidate genetic screen to identify natural genetic variation in the human ARHGEF26 gene that correlates with *Salmonella* invasion. Springboarding from this result, we experimentally tested and revised existing models of ARHGEF26’s role in *Salmonella* invasion, discovered an additional new role for ARHGEF26 during *Salmonella* disease, and confirmed our findings in mouse models. Building on how ARHGEF26 functions in other contexts, we implicated two ARHGEF26-interacting host proteins as contributors to *Salmonella* pathobiology. Collectively, these results identify a potential source of inter-person diversity in susceptibility to Salmonella disease, expand our molecular understanding of Salmonella infection to include a multifaceted role for ARHGEF26, and identify several important future directions that will be important to understand how *Salmonella* recruit and manipulate ARHGEF26 as well as how ARHGEF26 is able to drive *Salmonella*-beneficial processes.

## Introduction

The ability for bacteria to invade non-phagocytic host cells has long been recognized as a crucial trait of many pathogenic bacteria. Observations of this phenomena in *Salmonella* stretch back to at least 1920 when Margaret Reed Lewis observed that *Salmonella enterica* serovar Typhi (*S*. Typhi) induces vacuole formation during invasion of chick embryo tissues (1). With the advent of molecular biology, Galán and others identified that the type-III secretion system coded by genes in the *Salmonella* Pathogenicity Island-1 (SPI-1) facilitates *Salmonella* invasion (2, 3). Additional work demonstrated that the *Salmonella* effector proteins SopB, SopE, and SopE2 drive uptake of *Salmonella* in cultured cells by macropinocytosis through their ability to hijack or mimic host proteins (4–9). The importance of *Salmonella* invasion has been affirmed through several *in vivo* studies, as strains defective for the invasion apparatus are severely attenuated in their ability to colonize and disseminate (3), and/or drive inflammation (10, 11) in mouse models.

While much is known about the molecular mechanisms of *host-Salmonella* interactions, significantly less is known about why individuals have different susceptibilities to *Salmonella* infection. For example, in one recent *S*. Typhi human challenge study, the amount of *S*. Typhi found in patient blood varied substantially (0.05-22.7 CFU/mL blood in control patients), and 23% of participants resisted Typhoid fever onset (12). To help fill this gap, genome-wide association studies (GWAS) of Typhoid fever and non-typhoidal *Salmonella* bacteremia have demonstrated the importance of the HLA-region (13) and immune signaling (14) in *Salmonella* susceptibility. We hypothesized that differences in susceptibility to SPI-1 effectors and *Salmonella* host cell invasion also regulate risk of *Salmonella* infection. In fact, using a novel cellular genome-wide association platform called Hi-HOST (15–17), we previously determined that SNPs that affect *VAC14* expression regulate susceptibility to *S*. Typhi invasion through regulation of plasma membrane cholesterol (18). This demonstrates the power of cellular GWAS to identify natural genetic variation in cellular traits and enhance our mechanistic understanding of variable disease susceptibility.

In this work, we leveraged current understanding of host factors manipulated by *Salmonella* to identify human genetic variation that regulates invasion. We identified a locus in the guanine exchange factor (GEF) *ARHGEF26* (also known as *SGEF*) that correlated with susceptibility to *S*. Typhi and *S*. Typhimurium invasion. Previous work has demonstrated that ARHGEF26 contributes to *Salmonella*-induced membrane ruffling and was hypothesized to impact invasion (7). Current models speculate that ARHGEF26 contributes to membrane ruffling by enabling SopB-mediated activation of the human small GTPase RHOG. Here we demonstrated that ARHGEF26 regulates susceptibility to *Salmonella* invasion through *ARHGEF26* knockdown and overexpression. We also expanded our understanding of *Salmonella’s* interaction with ARHGEF26, finding that *S*. Typhi, but not *S*. Typhimurium uses ARHGEF26 for SopB- and SopE-mediated invasion of HeLa cells. Notably, we observed no effect of *RHOG* on invasion, in line with recent studies that found RHOG is dispensable for invasion (19, 20). In contrast, we show that reported interacting partners of ARHGEF26 in the Scribble complex, DLG1 (also known as SAP97) and SCRIB (also known as Scribble) also contribute to *S*. Typhi invasion. We demonstrated the importance of these findings *in vivo*, finding that *Arhgef26* deletion restricted *S*. Typhimurium burden in a mouse model of infection. Finally, we report a previously unappreciated role for ARHGEF26 in regulating the inflammatory response to *Salmonella* in both HeLa cells and mice. Collectively, these data identify a novel locus that contributes to natural genetic susceptibility to SPI-1-mediated invasion and elucidate our mechanistic understanding of ARHGEF26-mediated *Salmonella* invasion and inflammation.

## Results

### Cellular GWAS identifies an association between the rs993387 locus and *Salmonella* invasion

Our previous cellular GWAS (hereafter called H2P2; (17)) linked natural human genetic variation across 528 genotyped lymphoblastoid cell lines (LCLs) from parent-offspring trios to 79 infection phenotypes, including rates of *Salmonella* invasion. To quantify *Salmonella* invasion in H2P2, we infected LCLs with GFP-tagged *Salmonella enterica* serovar Typhi (*S*. Typhi) or *Salmonella enterica* serovar Typhimurium (*S*. Typhimurium) and counted the GFP^+^ host cells, which contain viable bacteria, by flow cytometry three hours post infection (Figure 1A). Using *Salmonella* invasion as a quantitative trait, we performed GWAS to identify loci associated with susceptibility to *Salmonella* invasion.

**Figure 1.**
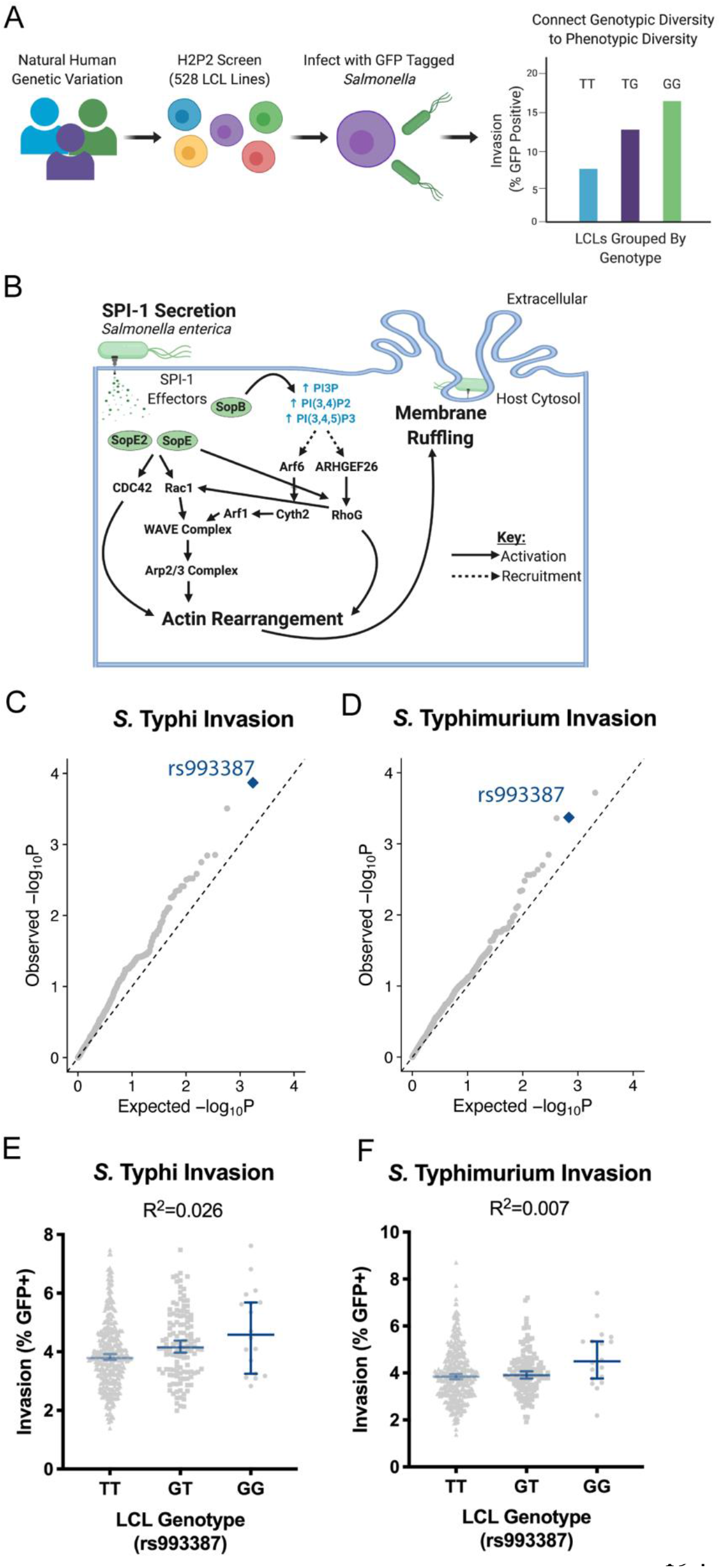
H2P2 reveals the rs993387 locus is associated with *Salmonella* invasion of lymphoblastoid cell lines. (A) Schematic for the H2P2 cellular GWAS. 528 lymphoblastoid cell lines (LCLs) from four populations were infected with *S*. Typhimurium (MOI 30) or *S*. Typhi (MOI 5) for 1 hour. Invasion was quantified 3 hours post infection by flow cytometry. Percent invasion was used as a phenotype for GWAS analysis. (B) Schematic for SPI-1-mediated invasion. Genes and complexes listed are included in the stratified GWAS analysis. (C,D) Stratified QQ plots examining SNPs associated with *S*. Typhi (C) and *S*. Typhimurium (D) invasion. Only SNPs in SPI-1 invasion-associated host genes were considered and analysis was restricted to common SNPs (MAF>0.05) and pruned at r^2^>0.6. rs993387 (blue diamond) diverges from p-values expected by chance for both serovars. Empirical P-values were calculated from family-based association analysis using QFAM-parents in PLINK. (E,F) Analysis of invasion for *S*. Typhi (E) and *S*. Typhimurium (F) from the H2P2 screen plotted by rs993387 genotype. Each dot represents a single LCL line, averaged between three independent experiments. Bar marks the median and the error bars represent the 95% confidence intervals. R^2^ values derived from simple linear regression. P values for both regressions were p≤0.05.

H2P2 identified 17 SNPs that passed genome-wide significance (p<5×10^−8^), however, no *Salmonella* invasion-associated SNPs passed this threshold (17). We next leveraged the last twenty years of *Salmonella* cellular microbiology and restricted our search space to common SNPs (minor allele frequency > 0.05) in 25 genes that regulate *Salmonella*-induced actin rearrangement, membrane ruffling, and/or invasion (Figure 1B, Table S1). These host genes encode proteins affected by SPI-1 secreted proteins (reviewed (21, 22)), and include *ARF1* (23), *ARF6* (23, 24)*, ARHGEF26* (commonly called *SGEF*) (7)*, RHOG* (7, 25)*, CYTH2* (23, 24), *CDC42* (7, 26–28), *RAC1* (7, 26, 28, 29), and actin (*ACTB*) (30–33), as well as genes in the WAVE (23, 28, 34) and Arp2/3 (27, 28, 34) complexes. Together, the proteins in this cascade lead to actin cytoskeletal rearrangements that enable macropinocytosis and bacterial uptake.

Plotting this SNP subset on a QQ plot to compare expected and observed p-values revealed a deviation towards p-values lower than expected by chance for both *S*. Typhi (Figure 1C) and *S*. Typhimurium (Figure 1D) invasion. The lowest p-value SNP associated with invasion of *S*. Typhi was rs993387 (p=0.0001), which is located in an *ARHGEF26* intron. This SNP also showed deviation with *S*. Typhimurium invasion (p=0.0004), although a linked SNP, rs71744878 (LD r^2^ =0.63 for ESN; 0.48 for GWD; 0.97 for KHV; 0.75 for IBS from LD Link (35)) had a slightly lower p-value. Removing all *ARHGEF26* SNPs from the analysis returned the remaining SNPs to the expected neutral distribution, suggesting, surprisingly, we only detect natural genetic variation in *ARHGEF26* that substantially impacted *Salmonella* invasion (Figure S1A, S1B). The rs993387 G allele associated with susceptibility to invasion, and the SNP appears to have a larger effect on *S*. Typhi invasion (2.6%, Figure 1E) compared to *S*. Typhimurium invasion (0.7%, Figure 1F). Within each of the four populations used in H2P2, the rarity of the minor allele made it difficult to comment on the impact of the GG genotype, but the directionality of effect between the TT and GT genotypes was preserved across all populations (Figure S1C).

### Manipulating *ARHGEF26* expression phenocopies the rs993387 locus’ effect on *Salmonella* invasion

We next analyzed the rs993387 locus in detail and found that SNPs in a ~100kb region of linkage disequilibrium overlapping *ARHGEF26* were associated with both *S*. Typhi (Figure 2A) and *S*. Typhimurium (Figure 2B) invasion. Looking for plausible functional variants in high LD (r^2^ > 0.6 in EUR and AFR populations) in Haploreg (36) revealed only additional intronic variants. Further evaluation of published eQTL datasets (37, 38) did not reveal a definitive connection to *ARHGEF26* mRNA expression. Testing for enhancer activity of a ~5kb region including rs993387, exon 11, and rs2122363 (the lowest *S*. Typhimurium SNP (Figure 2B)) using a luciferase reporter plasmid (39) demonstrated roughly two-fold induction over the vector control but no allele-specific enhancer activity in HeLa cells (Figure S1D). Additional analysis of the GTEx database (37) revealed that rs993387 is a splicing QTL in multiple tissues, including the colon (p=1.1×10^−9^), representing a plausible mechanism by which this SNP could regulate *ARHGEF26* protein abundance or function. In summary, while H2P2 implicated the *ARHGEF26* region in regulating susceptibility to *Salmonella* invasion, we do not yet know how genetic variation in this region affects *ARHGEF26* expression and/or function.

**Figure 2.**
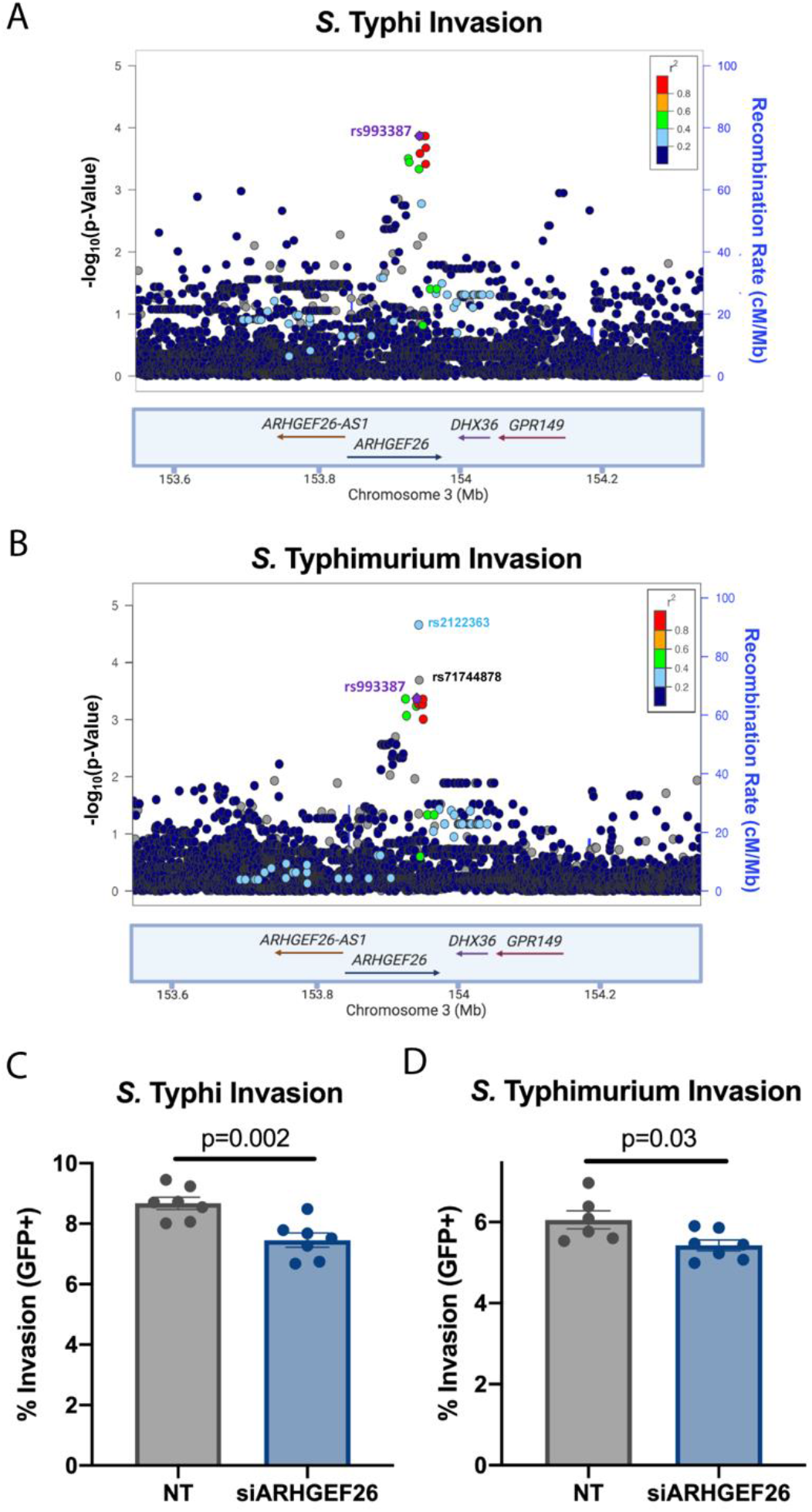
Knockdown of ARHGEF26 phenocopies human genetic variation in the rs993387 locus. (A,B) LocusZoom (41) plot generated with H2P2 data show SNPs in linkage disequilibrium with rs993387 in the *ARHGEF26* gene associates with *S*. Typhi (A) and *S*. Typhimurium (B) invasion. Height of dots represent the −log10(p-value) from H2P2. Dot color represents linkage disequilibrium (r^2^) based on 1000 Genomes African dataset. Blue line behind dots tracks the recombination rate. (C,D) RNAi-mediated ARHGEF26 knock down reduces *S*. Typhi (siARHGEF26/NT = 0.86, C) and *S*. Typhimurium (siARHGEF26/NT = 0.90, D) invasion in LCLs (HG01697, IBS, rs993387 genotype=GG) compared to non-targeting (NT) siRNA. Cells were infected at MOI 5 (*S*. Typhi) or MOI 30 (*S*. Typhimurium) for 60 minutes. Invasion was measured three hours post infection by flow cytometry. Each dot represents a biological replicate from one of three independent experiments. Experimental means were adjusted to the grand mean prior to plotting or performing statistics. Bars represent the mean and error bars represent standard error of the mean. P-values generated by an unpaired t-test.

We next examined if *ARHGEF26* expression affects *Salmonella* invasion in LCLs. While previous reports have linked *ARHGEF26* to the induction of membrane ruffling (7), no study has demonstrated whether *ARHGEF26* contributes to host-cell invasion. This is an important distinction, as invasion does not always correlate with ruffling (40). RNAi knockdown of *ARHGEF26*, confirmed by qPCR (Figure S1E), showed reduced *S*. Typhi and *S*. Typhimurium invasion—a phenotype similar to the protective rs993387 T-allele in LCLs (Figure 2C, D).

### ARHGEF26 effects on invasion are cell line and serovar dependent

To dissect how ARHGEF26 contributes to *Salmonella* invasion, we examined how the protein regulates *Salmonella* invasion of HeLa cells, a common epithelial invasion model. We hypothesized that our core phenotypes of *ARHGEF26* positively regulating *S*. Typhi and *S*. Typhimurium invasion in LCLs would replicate in HeLa cells. To our surprise, *ARHGEF26* RNAi knockdown significantly reduced *S*. Typhi invasion into HeLa cells (Figure 3A) but did not affect *S*. Typhimurium invasion (Figure 3B). This revealed that invasion only depends on *ARHGEF26* in certain cell line and serovar combinations.

**Figure 3.**
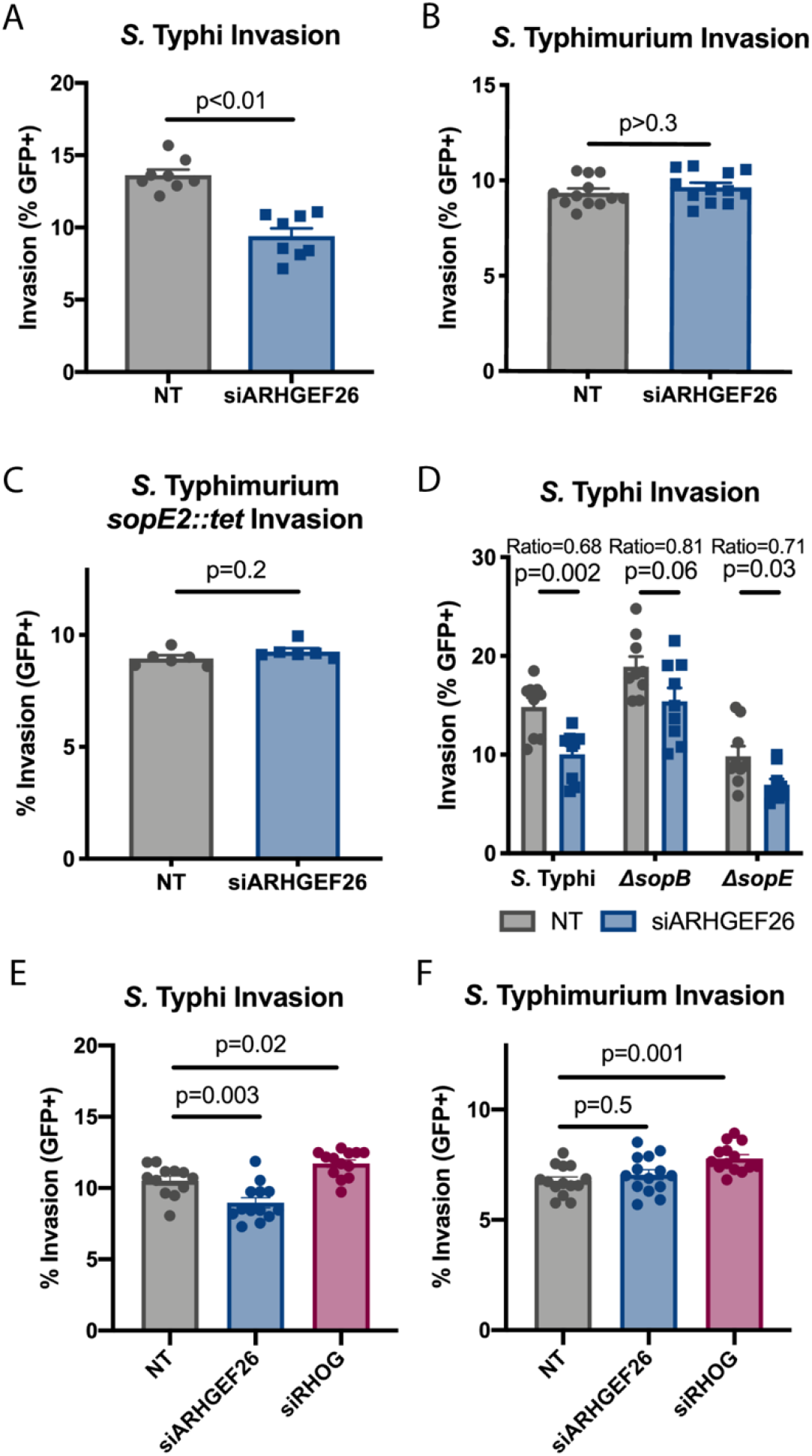
*ARHGEF26* is a positive regulator of *Salmonella* Typhi, but not *Salmonella* Typhimurium, invasion into HeLa cells. (A,B) RNAi knockdown of ARHGEF26 in HeLa cells results in reduced *S*. Typhi (A), but not reduced *S*. Typhimurium (B) invasion. (C) The absence of an effect in *S*. Typhimurium is independent of *sopE2*, as a *S*. Typhimurium strain where the gene is replaced with a tetracycline resistance allele (*tet*) also shows no effect. (D) The effects of *ARHGEF26* knockdown on *S*. Typhi invasion does not require either *sopB* or *sopE*. (E, F) *RHOG* knockdown does not reduce *S*. Typhi or *S*. Typhimurium invasion. All comparisons are made to transfection with non-targeting (NT) siRNA. Cells were infected at MOI 30 (*S*. Typhi) or MOI 1 (*S*. Typhimurium) for 30 minutes. For all panels, invasion was measured three hours post infection by flow cytometry. All dots represent biological replicates from at least two experiments. Ratio in D is the siARHGEF26 mean divided by the NT mean. Data across experiments were normalized to the grand mean prior to plotting or performing statistics. Bars represent the mean and error bars represent standard error of the mean. P-values for panels A-D were generated by unpaired t-test. P-values for E-F were generated by one-way ANOVA with Dunnett’s multiple comparisons test.

The prevailing model for *ARHGEF26* involvement in invasion is that Salmonellae use SopB to recruit ARHGEF26 and thereby activate RHOG, while SopE and SopE2 independently and directly activate RHOG (7). Based on this, we speculated that SopE2 (present in *S*. Typhimurium 14028s; absent in *S*. Typhi Ty2) might be a more potent activator of RHOG in HeLa cells than SopE (present in *S*. Typhi Ty2; absent in *S*. Typhimurium 14028s), and that this effector repertoire difference might explain our difference in *ARHGEF26*-dependent invasion. However, in an *S*. Typhimurium strain lacking SopE2 (leaving only SopB to drive invasion) there was no effect of *ARHGEF26* knockdown on invasion (Figure 3C). Notably, the *S*. Typhimurium 14028s and *S*. Typhi Ty2 SopB proteins are 98.4% identical on the amino acid level, so it is possible that the modest differences in sequence could account for the differential ARHGEF26 requirement. A potentially more plausible hypothesis is that additional differences in the effector repertoires and/or invasion mechanisms of *S*. Typhi and *S*. Typhimurium enable *S*. Typhimurium to efficiently invade HeLa cells in the absence of *ARHGEF26*.

### ARHGEF26 contributes to SopB- and SopE-mediated *Salmonella* invasion

We next tested the hypothesis that ARHGEF26 is required for SopB-mediated invasion, but not SopE-mediated invasion using Δ*sopB* and Δ*sopE S*. Typhi. Surprisingly, *ARHGEF26* knockdown reduced both SopB- and SopE-mediated invasion (Figure 3D). With wild-type *S*. Typhi, we observe a robust reduction in invasion following *ARHGEF26* knockdown (siARHGEF26/NT = 0.68). We observed a more modest reduction in invasion following *ARHGEF26* knockdown with the Δ*sopB* mutant (siARHGEF26/NT = 0.81), demonstrating that SopE-mediated invasion is less efficient without ARHGEF26. In contrast, with the Δ*sopE* mutant, we saw almost no change in effect size (siARHGEF26/NT = 0.71), demonstrating that when only SopB is present, ARHGEF26 has roughly the same proportional effect on invasion as when both effectors are present. Together, these data suggest that the previous model in which ARHGEF26 is recruited by only SopB is incomplete, and instead support a model in which ARHGEF26 contributes to both SopB- and SopE-mediated invasion.

### *RHOG* knockdown does not phenocopy *ARHGEF26* knockdown

We next tested whether the effects of *ARHGEF26* knockdown could be phenocopied by knocking down *RHOG*. Notably while Patel *et al*. showed that RHOG contributes to SopB-mediated membrane ruffling (7) and invasion (25), more recent reports have demonstrated that RHOG is dispensable for *Salmonella* invasion into fibroblasts (20) and Henle cells (19, 20). In line with this, we found that *RHOG* knockdown did not reduce *Salmonella* Typhi (Figure 3E) or Typhimurium (Figure 3F) invasion. Curiously, we found that *RHOG* knockdown actually subtly increased invasion.

We hypothesize that there are three non-mutually exclusive reasons why *RHOG* knockdown fails to phenocopy *ARHGEF26* knockdown. First, it is possible that 85% knockdown (Figure S1F) may not be sufficient to impair RHOG’s cellular functions, particularly in light of the small fraction of active RHOG in the cell at any given time (42–44). Alternatively, ARHGEF26-mediated invasion may involve other small GTPases. For instance, ARHGEF26 does show a weak capacity to stimulate nucleotide exchange of CDC42 and RAC1 *in vitro* (45, 46). Alternatively, ARHGEF26 could activate other CDC42-related small-GTPases, such as RHOJ, which have recently been shown to affect invasion (19, 47). Our third hypothesis is that ARHGEF26 may have impacts on *Salmonella* invasion independent of its GEF activity. This is based on work from one of our labs that demonstrated that some of ARHGEF26’s cellular roles involve forming a tertiary complex with DLG1 and SCRIB independent of nucleotide exchange (48).

### The Scribble Complex members DLG1 and SCRIB promote *S*. Typhi invasion

Previous work has demonstrated that the Scribble complex members DLG1 and/or SCRIB play key roles facilitating ARHGEF26 activity during human papillomavirus infection (49), as well as in regulating epithelial junctions formation, contractability, and lumen formation in 3D cysts (48). With this in mind, we investigated whether DLG1 and/or SCRIB has a role in *Salmonella* invasion. Indeed, we observed that knockdown of *DLG1* and *SCRIB* each resulted in a reduction in *S*. Typhi invasion (Figure 4A).

**Figure 4:**
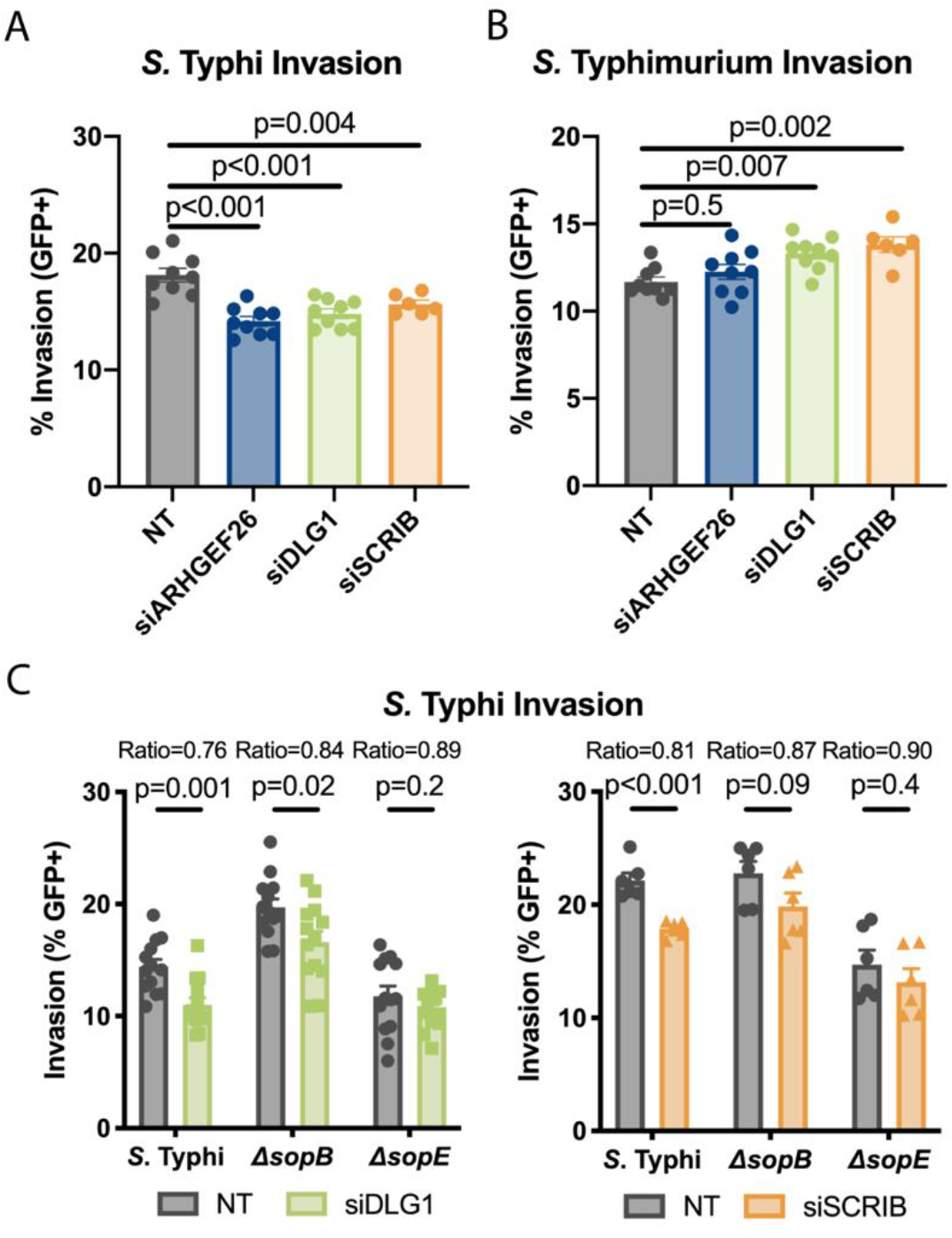
ARHGEF26 interactors DLG1 and SCRIB contribute to *S*. Typhi invasion. (A,B) RNAi knockdown of *DLG1* and *SCRIB* phenocopy the reduction in *S*. Typhi (A), but not *S*. Typhimurium (B) invasion that we observe with *ARHGEF26* knockdown. (C) *DLG1* and *SCRIB* knockdown significantly reduces wild-type *S*. Typhi and Δ*sopB S*. Typhi invasion, but not Δ*sopE* invasion. Cells were infected at MOI 30 (*S*. Typhi) or MOI 1 (*S*. Typhimurium) for 30 minutes. Circles represent non-targeting (NT) siRNA, squares siDLG1, and triangles siSCRIB treated wells. Invasion was measured three hours post infection by flow cytometry. All dots represent biological replicates from at least three experiments. Ratio in C represent the mean of invasion of siRNA treatment divided by the mean invasion of NT treatment. Data across experiments were normalized to the grand mean prior to plotting or performing statistics. Bars represent the mean and error bars represent standard error of the mean. P-values were generated by unpaired one-way ANOVA with Dunnett’s multiple comparison test for A and B. For panel C p-values were generated by unpaired t tests.

Supporting our hypothesis that DLG1, SCRIB, and ARHGEF26 may act through the same pathway, our findings with *DLG1* and *SCRIB* knockdown broadly phenocopy our results with *ARHGEF26* knockdown. Consistent with our *ARHGEF26* knockdown data, *DLG1* and *SCRIB* are dispensable for *S*. Typhimurium invasion (Figure 4B). Further, the effect of *DLG1* and *SCRIB* knockdown on invasion was partially reduced when cells were infected with Δ*sopB S*. Typhi (Figure 4C, siDLG1/NT = 0.84, p=0.02; siSCRIB/NT = 0.87, p =0.09) compared to infection with wild-type *S*. Typhi (Figure 4C, siDLG1/NT = 0.76, p = 0.001; siSCRIB/NT = 0.81, p < 0.001), suggesting that SopE is more efficient at inducing invasion in the presence of ARHGEF26, DLG1, and SCRIB. However, unlike *ARHGEF26* knockdown, the effect of *DLG1* and *SCRIB* knockdown were largely ablated when cells were infected with Δ*sopE S*. Typhi (Figure 4C, siDLG1/NT = 0.89; siSCRIB/NT = 0.90), demonstrating that DLG1 and SCRIB have small and statistically insignificant impacts on SopB-mediated invasion. Our interpretation of this data is that DLG1 and SCRIB may primarily help ARHGEF26 drive SopE-mediated invasion, but that other scaffolds primarily assist in SopB-mediated invasion.

### The impacts of ARHGEF26 on *S*. Typhi invasion require GEF catalytic activity

Involvement of DLG1 and SCRIB in ARHGEF26-mediated *Salmonella* invasion could be facilitated by one of two mechanisms. First, DLG1 and SCRIB could help ARHGEF26 localize to the site of invasion in order to serve as a GEF, similar to how they function to drive adherens junction formation (48). Alternatively, formation of the ARHGEF26, DLG1, SCRIB tertiary complex could drive *Salmonella* invasion through unknown and GEF-independent mechanisms, as one of our labs described for actomyosin contractility (48). To distinguish between these possibilities, we assessed which ARHGEF26 domains are required for *S*. Typhi invasion by structure-function analysis (Figure 5A).

**Figure 5:**
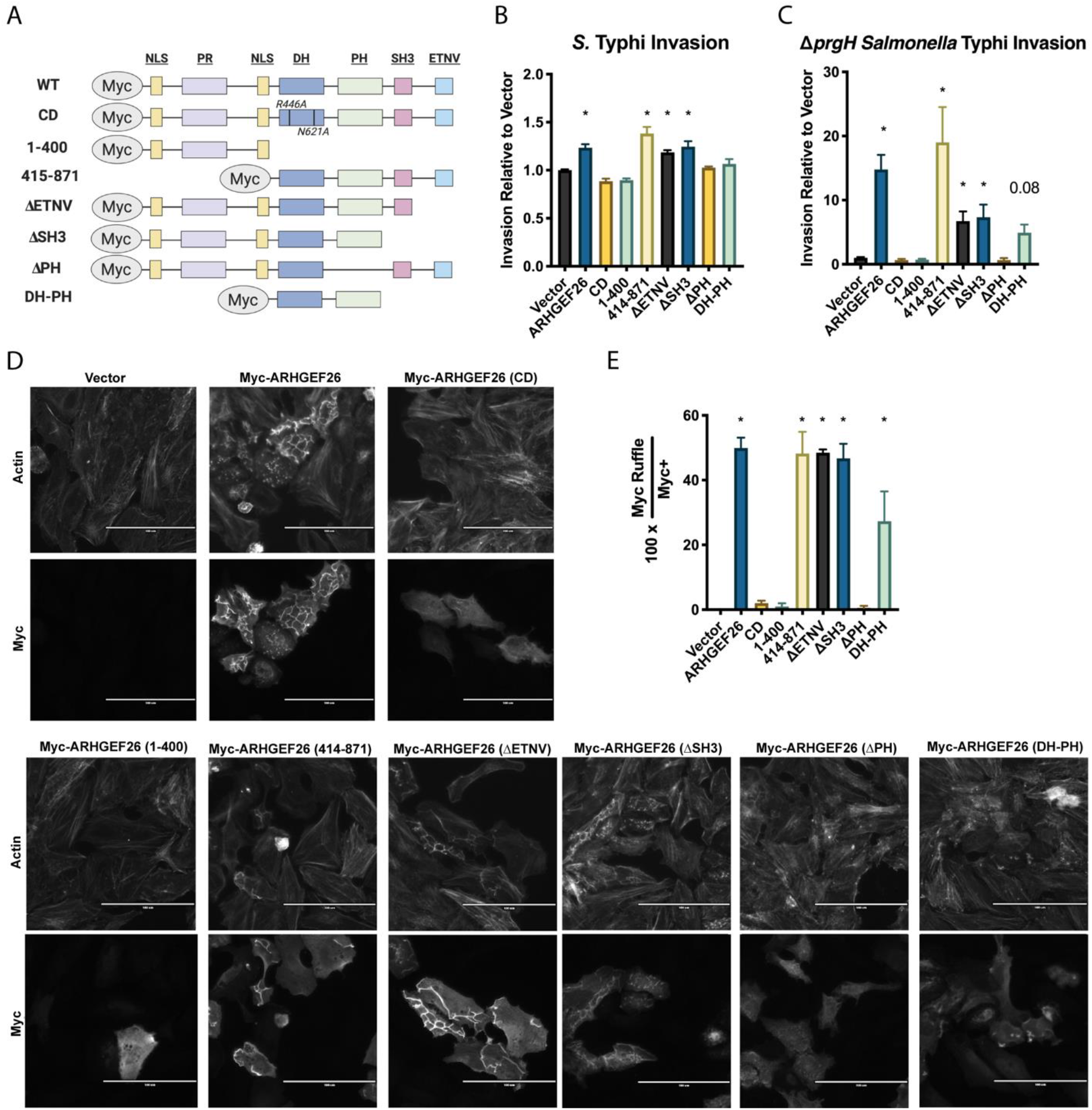
ARHGEF26 DH and PH domains are required for ARHEGEF26 to induce membrane ruffling and *Salmonella* invasion in HeLa cells. (A) Schematic of overexpression constructs used. (B,C) Overexpression of *ARHGEF26* constructs in HeLa cells results in increased wild-type *S*. Typhi invasion (B), as well as invasion of the SPI-1 secretion mutant Δ*prgH S*. Typhi (C). Infections were performed at MOI 30 for 60 minutes and quantified 3 hours post infection by flow cytometry. Invasion is reported relative to vector and includes data from at least three independent experiments with three replicates per experiment. * represents a corrected p-value < 0.05. P-values generated by Kruskal-Wallis test with Dunn’s multiple comparison test. (D) *ARHGEF26* constructs able to induce invasion also induce membrane ruffling. Overexpression of catalytically active constructs in HeLas results in membrane ruffling that can be observed both with ARHGEF26 staining using the Myc Tag, as well as by using phalloidin staining to observe actin. (E) Quantification of membrane ruffling confirms correlation with *ARHGEF26*-mediated invasion. Frequency of membrane ruffles was quantified as a percent of Myc+ cells that had a clear Myc+ ruffle. Ruffle abundance was quantified from four independent experiments. Values were generated by quantifying ruffles from five separate fields of view. The presence of a ruffle was confirmed by examining phalloidin stained actin at that site. Scale bar is 100 μM. * represents a p-value < 0.05. p-value generated by one-way ANOVA of the log(X+1) transformed values with Dunnett’s multiple comparisons test.

Overexpression of ARHGEF26, but not overexpression of a catalytically dead (CD) mutant (R446A, N621A), increased *S*. Typhi invasion (Figure 5B). Additionally, overexpression of mutants lacking the pleckstrin homology (PH) and/or the catalytic Dbl homology (DH) domain failed to induce invasion. All other domains were independently dispensable. While the DH and PH domains were necessary, they were not sufficient, as a DH-PH construct failed to induce invasion.

As catalytically active *ARHGEF26* overexpression induces spontaneous membrane ruffling and macropinocytosis, we next examined whether membrane ruffling and invasion could be decoupled. To do this, we overexpressed the *ARHGEF26* constructs and quantified the uptake of passive Δ*prgH S*. Typhi, which cannot dock to host cells or induce invasion (Figure 5C), as well as the number of ARHGEF26^+^ membrane ruffles present (Figure 5D, 5E). Across five of our six *ARHGEF26* mutants, we found strong correlations between *S*. Typhi invasion (Figure 5B), Δ*prgH S*. Typhi invasion (Figure 5C), and membrane ruffling (Figure 5D, 5E). The one exception was our PH-DH mutant, which was not able to promote wild-type *S*. Typhi invasion but did modestly increase Δ*prgH S*. Typhi invasion (p=0.08) and induce small membrane ruffles. This indicates that the small ruffles are sufficient to drive Δ*prgH* uptake but are an insignificant addition to the typical *S*. Typhi induced membrane ruffles. Overall, these data suggest that *ARHGEF26* overexpression affects invasion by increasing membrane ruffling and macropinocytosis.

We conclude that, under overexpression conditions, *ARHGEF26* constructs that do not show catalytic activity but are still able to bind DLG1 and SCRIB (48) are not able to drive *Salmonella* invasion. Thus, our RNAi and overexpression results support a model where both interaction of DLG1/SCRIB and nucleotide exchange contribute to ARHGEF26’s role in promoting invasion.

### ARHGEF26 does not show significant phosphoinositide binding using a dot blot assay

While our data suggest that DLG1 and SCRIB may contribute to ARHGEF26 localization, the prevailing model postulates that ARHGEF26 is guided to the plasma membrane through its pleckstrin homology (PH) domain (7), which in some proteins can bind phosphoinositides (50). Under this model, ARHGEF26 localization is regulated by SopB’s effects on phosphoinositides (6, 7, 9, 21, 51–53). Supporting this, our data demonstrates that the ARHGEF26 PH domain is required to increase invasion (Figure 5). However, many PH domains either completely lack canonical phosphoinositide binding or have phosphoinositide binding that is physiologically irrelevant (50, 54). Therefore, PH-dependence alone is insufficient to implicate SopB-generated phosphoinositides as this could suggest either that ARHGEF26 binds phosphoinositides in order to function, or simply, that this mutation disrupts the catalytic domain as has been shown for other RHO GEFs (46).

To directly test whether ARHGEF26 binds phosphoinositides, we performed a dot blot assay in which different phosphoinositide species are dotted on a membrane and exposed to ARHGEF26. Across a variety of conditions—including different ARHGEF26 constructs, cellular sources of protein, and blocking solutions—we did not detect strong phosphoinositide binding (Figure S2). We also tested whether co-expression of ARHGEF26 and RHOG could drive phosphoinositide binding, as occurs with the RHOG GEF Trio (55), but did not observe increased signal. Under some conditions, weak and non-specific signal appeared on the dots of some phosphoinositide species, but this was independent of the PH domain and difficult to distinguish from background noise. This contrasted with our positive control, the AKT-PH domain, which demonstrated robust and highly specific binding. By considering the difference in signal between the ARHGEF26 constructs and AKT-PH domain, as well as the history of this assay exaggerating the affinity for proteins with phosphoinositides (56), we surmise that even if this signal is the result of a weak affinity for phosphoinositides, this affinity is unlikely to play any physiological role *in vivo*. Therefore, while we cannot firmly rule out that ARHGEF26 binds phosphoinositides using this assay, these data do not support PH domain-mediated phosphoinositide binding directing ARHGEF26 localization.

Together, our results suggest a new model for the role of *ARHGEF26* during *S*. Typhi epithelial cell invasion. We propose that ARHGEF26 potentially stimulates invasion independently of RHOG and that recruitment of ARHGEF26 to the site of invasion is not dependent on SopB-mediated phosphoinositide changes. Instead, our results demonstrate the SCRIB-DLG1-ARHGEF26 complex is important for invasion even with SopE stimulation being the primary route of invasion. Our results that *ARHGEF26*, *SCRIB*, and *DLG1* knockdown have their strongest effects when both SopB and SopE are present may suggest these effectors must work cooperatively to effectively enable ARHGEF26 to enhance invasion.

### ARHGEF26 contributes to *S*. Typhimurium-induced inflammation in HeLa cells

In addition to enabling *Salmonella* invasion, interactions between the SPI-1 secreted effectors and host machinery drive inflammation that is characteristic of *Salmonella* infections. For instance, during *Salmonella* invasion, SopB and SopE activate CDC42, which goes on to enable the formation of a PAK1–TRAF6-TAK1 complex, NF-κB activation, and increased IL-8 production (7, 57–61). Other work has suggested NOD2 and RIPK2 also contribute to CDC42 and RAC1 mediated inflammation (62). Further, studies using dominant negative constructs have suggested that CDC42 and Rac1 are required for secretion of IL-8 from polarized cells *in vitro*, presumably through changes to the cytoskeleton (63). Based on our findings that ARHGEF26 is a critical GEF during *Salmonella* invasion, we hypothesized that it may also have a role in mediating *Salmonella-induced* inflammation.

To examine how ARHGEF26 contributes to inflammation, we knocked down *ARHGEF26* and *RHOG* in HeLa cells and measured IL-8 abundance in supernatant. In supernatant from uninfected cells, IL-8 was reduced following *ARHGEF26* or, interestingly, *RHOG* knockdown (Figure 6A). This suggests that even under basal conditions, ARHGEF26 regulates inflammation, potentially through interactions with RHOG.

**Figure 6:**
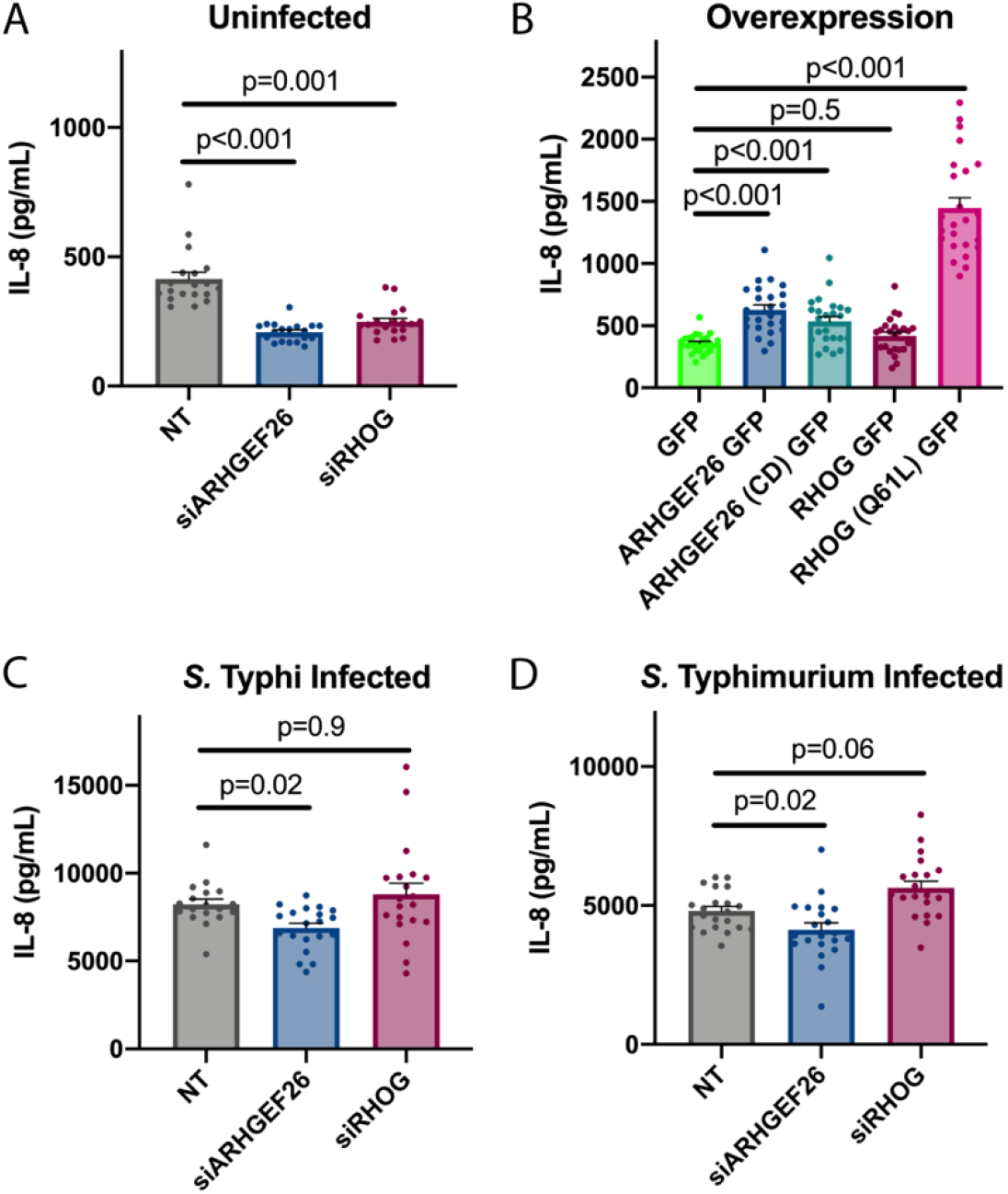
ARHGE26 and RHOG are context-dependent enhancers of IL-8 abundance in HeLa cell supernatant. (A) *ARHGEF26* and *RHOG* knockdown results in less IL-8 secretion into HeLa supernatant than non-targeting (NT) siRNA. Media was changed two days post transfection supernatant was collected 8 hours later. (B) Overexpression of *ARHGEF26* and *RHOG* in HeLa cells increases IL-8 cytokine abundance in supernatant. Media was changed on transfected cells 18-24 hours post transfection and supernatant was collected 6 hours later. (C, D) *ARHGEF26*, but not *RHOG*, knockdown reduces IL-8 abundance in supernatant following *S*. Typhi infection (C) and *S*. Typhimurium infection (D). For C and D, cells were infected two days post transfection. Cells were infected with *S*. Typhi (MOI 30) or *S*. Typhimurium (MOI 1) for one hour before gentamycin addition. Media was changed two hours before infection and supernatant was collected six hours after infection. Cytokine abundance was measured by ELISA. For all graphs, dots represent a single well and data were collected across seven independent experiments. Data were normalized to the grand mean prior to plotting or performing statistics. For (C), two outliers identified by ROUT (Q=0.1%) were removed from the non-targeting group. These values (17,924 pg/mL and 22,083 pg/mL) inflated the mean of the NT group, making the ARHGEF26 effect size artificially large, and the p-value artificially low (p=0.002). P-values were calculated using a one-way ANOVA with Dunnett’s multiple comparison test on the log_2_ transformed data. For all graphs central tendency is the mean, error bars are the standard error of the mean.

Overexpression experiments confirmed the importance of ARHGEF26 and RHOG in regulating basal inflammatory cytokine production. *ARHGEF26* overexpression resulted in significantly increased IL-8 production (Figure 6B). Surprisingly, a partial effect was observed with overexpression of the catalytically dead construct (Figure 6B), demonstrating that ARHGEF26 does not require GEF activity to influence cytokine production. Overexpression of wild-type *RHOG* did not increase IL-8 in supernatant, but overexpression of a constitutively active *RHOG* construct resulted in very robust cytokine production (Figure 6B). Together, our knockdown and overexpression data demonstrate that ARHGEF26 promotes inflammatory cytokine production and involves both GEF-dependent and GEF-independent mechanisms.

We next sought to examine whether ARHGEF26 regulates inflammation during *Salmonella* infection. Infection with *S*. Typhi caused induction of IL-8, but levels were moderately lower in supernatants with *ARHGEF26*, but not with *RHOG* knockdown (Figure 6C). This aligned with what we observed with invasion, suggesting either that related mechanisms could be contributing to ARHGEF26-mediated invasion and inflammation, or that reduced inflammation is driven by reductions in invasion. However, in contrast to our invasion data, we found that *ARHGEF26*, but not *RHOG*, knockdown also moderately reduced IL-8 abundance following *S*. Typhimurium infection (Figure 6D). This suggests that *S*. Typhimurium-mediated ARHGEF26-enhanced IL-8 production is invasion independent. Together, these data demonstrate that ARHGEF26 and RHOG have context specific roles through which they promote inflammation in HeLa cells.

### ARHGEF26 is required for *S*. Typhimurium virulence during an enteric fever model of infection

Following these results, it was crucial to define ARHGEF26’s role in *Salmonella* pathogenesis in mice. As *S*. Typhi is a human specific pathogen, we focused specifically on how ARHGEF26 influences murine *S*. Typhimurium pathogenesis. Based on the data from H2P2 and LCL knockdown, we hypothesized that ARHGEF26 is required for SPI-1 mediated establishment of *S*. Typhimurium in the mammalian gut. To test this hypothesis, we utilized *Arhgef26^-/-^* C57BL/6J mice (64) to assess the ability of *S*. Typhimurium to establish infection using the oral enteric fever model of infection (Figure 7A). In this model, SPI-1 secretion is required for *S*. Typhimurium colonization and persistence in the mammalian ileum and helps facilitate *S*. Typhimurium dissemination to the spleen (3). In support of our hypothesis that *Arhgef26* promotes invasion *in vivo, Arhgef26^-/-^* mice showed significantly lower *S*. Typhimurium burdens in the ileum, but this effect was considerably smaller in the spleen (Figure 7B).

**Figure 7.**
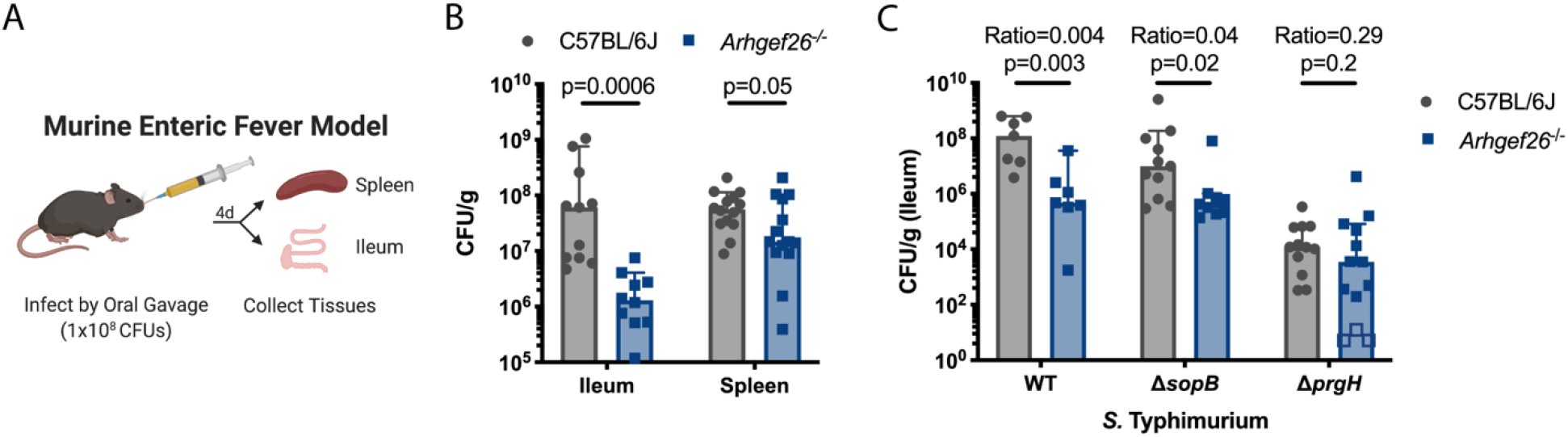
*Arhgef26* is critical during *S*. Typhimurium infection in the enteric fever murine model. (A) Schematic of the murine enteric fever infection model. (B) *Arhgef26^-/-^* mice have reduced ileal and splenic bacterial burden in the enteric fever infection model compared to C57BL/6J mice. P-values generated by two-way ANOVA with Sidak’s multiple comparison test. (C) Effects of *Arhgef26* knockout on bacterial fitness depends on *sopB* and *prgH*. Open boxes represent mice where no colony forming units (CFU) were recovered from *ARHGEF26^-/-^* mice and the CFU/g was set to the limit of detection. Ratio represents median CFU/g recovered from *Arhgef26^-/-^* mice divided by the median CFU/g recovered from C57BL/6J mice. P-values were generated by unpaired t-tests. For all experiments, mice were infected with 1×10^8^ *S*. Typhimurium CFU and tissues were harvested four days post infection for CFU quantification. For all bar graphs, each dot represents a single mouse from one of at least two experiments, bars represent the median, and error bars represent the 95% confidence interval. All mice were age and sex matched within experiments, with both sexes represented in all experiments.

As *Arhgef26^-/-^* mice phenocopy the effects of SPI-1 knockout on ileal burden, we next examined whether the effect of *Arhgef26* knockout requires SPI-1. To genetically test this hypothesis, wild-type and *Arhgef26^-/-^* mice were infected with wild-type, Δ*sopB*, and Δ*prgH S*. Typhimurium. Supporting that a functional SPI-1 secretion system is required for ARHGEF26 to affect *S*. Typhimurium fitness, the difference in burden between wild-type and *Arhgef26^-/-^* mice was significantly reduced when mice were infected with Δ*prgH* bacteria (Figure 7C). Notably, a few *Arhgef26* knockout mice did show Δ*prgH* ileal burdens below the limit of detection, possibly suggesting an additional yet inconsistent level of resistance in these mice. This could be the result of reduced uptake of bacteria by phagocytes, as a previous screen reported that *ARHGEF26* is required for *Salmonella* uptake by macrophages (65). Interestingly, we also observe a reduced effect of *Arhgef26* deletion on bacterial fitness with the Δ*sopB* mutant (Figure 7C). Interestingly, this is reminiscent of the smaller phenotype of Δ*sopB S*. Typhi with *ARHGEF26* RNAi in HeLa cells.

### ARHGEF26 contributes to inflammation during a gastroenteritis model of *S*. Typhimurium infection

While SPI-1 contributes significantly to the colonization and survival of *S*. Typhimurium in the ileum in the enteric fever model of infection, this is not the natural progression of most *S*. Typhimurium illness in humans. Instead, *S*. Typhimurium typically causes severe gastroenteritis and diarrheal disease, with notable exceptions (66). To model this natural disease progression in mice, microbiota must be reduced with streptomycin pretreatment prior to infection (11) (Figure 8A). Interestingly, in this model, SPI-1 secretion does not impact bacterial burden at early timepoints but instead drives severe inflammation (10, 11). As we have demonstrated that ARHEGF26 is a regulator of the cytokine response (Figure 6), we examined whether ARHGEF26 promoted SPI-1 mediated pathology by infecting wild-type and *Arhgef26^-/-^* mice following streptomycin pretreatment. Indeed, there was no effect of *Arhgef26* deletion on the recovery of *S*. Typhimurium from ileum, cecum, or colon two days post infection (Figure 8B), but these sites demonstrated significantly reduced inflammation-associated pathology in *Arhgef26^-/-^* mice (Figure 8C, 8D). This was most striking in the cecum and colon where inflammation is most severe in wild-type mice. The data from both models demonstrate that ARHGEF26 plays a critical, multifaceted role in enabling *S*. Typhimurium to utilize the SPI-1 secretion system to cause disease during murine infection.

**Figure 8.**
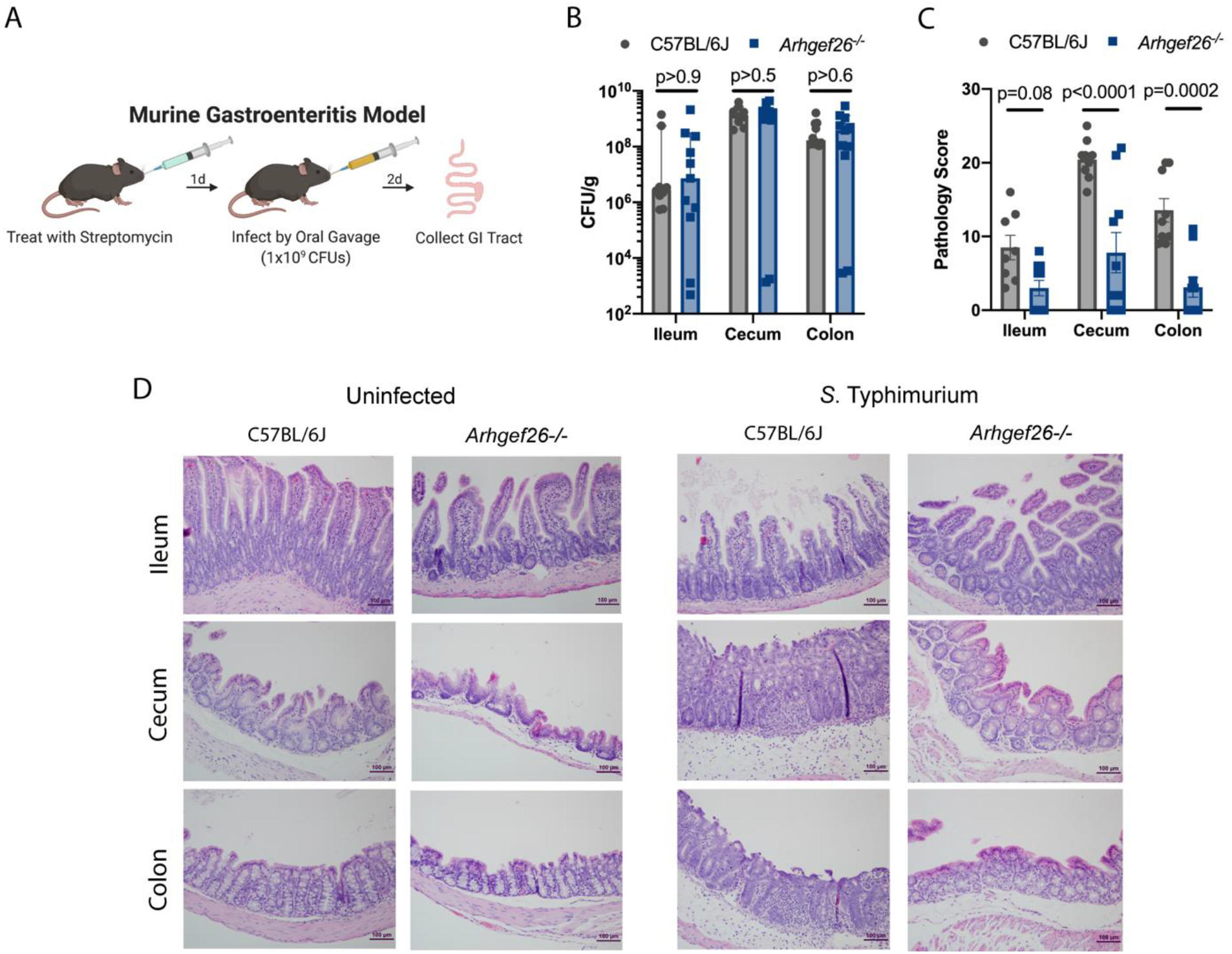
*Arhgef26* enhances inflammation in mice. (A) Schematic for murine gastroenteritis infection model. Mice are pretreated with streptomycin one day before infection with 1×10^9^ *S*. Typhimurium CFUs. Tissues are collected 2 days post infection for CFU quantification or histological analysis. (B) *Arhgef26^-/-^* mice have no reductions in gastrointestinal tract bacterial burdens in the gastroenteritis infection model. (C) *Arhgef26^-/-^* mice have significantly reduced inflammation following infection compared to C57BL/6J mice in the gastroenteritis infection model. Pathology scores were generated in a blinded fashion by a trained pathologist and are broken down in Supplemental File 1. For B and C, each dot represents a single mouse from one of two experiments, bars represent the median and error bars represent the 95% confidence interval. All mice were age- and sex-matched within and across experiments. P-Values generated by two-way ANOVA with Sidak’s multiple comparison test. (D) Examples of differential pathology following infection between C57BL/6J and *Arhgef26^-/-^* mice. Scale bar is 100 μM.

## Discussion

Beginning with a candidate pathway approach paired with the Hi-HOST cellular GWAS platform, we identified a QTL in the *ARHGEF26* gene that influences host cell invasion, carried out functional studies that reshape our understanding of how ARHGEF26 stimulates invasion, and revealed a new role for ARHGEF26 in regulating inflammation in cells and mice.

One important question moving forward is how the rs993387 locus contributes to *Salmonella* invasion. As noted, published eQTL datasets have not provided a consistent answer for the association of rs993387 with *ARHGEF26* mRNA levels (37). This inconsistency is likely driven by the fact that while ARHGEF26 expression is detectable based on quantitative PCR (CT ~30) and RNA-seq datasets in LCLs (38, 67), it is a low abundance transcript. Additionally, we have not been able to reliably detect protein levels using either antibodies or mass spectrometric approaches. Of note, there is an ARHGEF26 anti-sense transcript (*ARHGEF26-AS1*) whose expression is associated with rs993387 in some tissues (most strongly in tibial nerve p=6.9×10^−22^) (37). Also rs993387 is reported as an *ARHGEF26* splicing QTL in both sigmoid (p=1.1×10^−9^) and transverse colon (p=5.4×10^−5^) (37, 38). Thus, though there are several plausible mechanisms, we do not know if rs993387 (or a causal variant in LD) affects ARHGEF26 mRNA levels, splicing, protein levels, or protein function, and such experiments have been technically challenging.

A second unanswered question is how natural variation regulating *ARHGEF26* could impact invasion. Small changes in *ARHGEF26* expression or function could change the rate of membrane ruffling as we observe in the knockdown and overexpression systems. Alternatively, changes in *ARHGEF26* could impact the size of membrane ruffles and the efficiency with which macropinocytosis occurs at the site of invasion. A comparable phenomenon is observed with our overexpression system, as constructs with low activity (*e.g*. the DH-PH construct) form small, contained ruffles (Figure 5D). In contrast, other constructs (*e.g*. the 414-871 construct) form large ruffles that were able to increase *S*. Typhi and Δ*prgH S*. Typhi invasion to levels even above what we observe with wild-type *ARHGEF26* overexpression (Figure 5B, 5C). Changing the size of membrane ruffles could have impacts beyond simply enabling the ARHGEF26-recruiting bacteria to invade the cell, as previous work has demonstrated that *Salmonella* swimming at the cell surface use membrane ruffling induced by other bacteria as a signal to begin their own invasion, thus engaging in a sort of cooperative behavior (68). Thus, large ARHGEF26-induced membrane ruffles could serve as a mechanism for (a) efficient macropinocytosis, and (b) cooperative host cell invasion.

Our work has also expanded understanding of ARHGEF26-mediated invasion by demonstrating that the protein has serovar and cell line dependent roles in *Salmonella* invasion, and that it appears to contribute to both SopB- and SopE-mediated invasion. This may indicate a context-dependent role for ARHGEF26 and raises a number of technical and conceptual questions. At present, we are unsure why this serovar specific interaction occurs, but our data suggest that it is independent of SopE and SopE2. Further, that serovar specificity occurs in the canonical *Salmonella* invasion model (HeLas), but not in LCLs, raises additional questions broadly about how representative HeLa cells are for studying invasion. Supporting this, our murine data, in which *S*. Typhimurium had a SPI-1 dependent fitness deficit in *Arhgef26^-/-^* mice (Figure 7C), are more consistent with our results from H2P2 (Figure 1F) and siRNA in LCLs (Figure 2D) than our HeLa cell data (Figure 3B). This reinforces a recent observation that there are striking differences in the invasion mechanisms observed in canonical tissue culture models compared to those observed *in vivo* (69, 70). These data serve as a reminder that host-pathogen biology is more complex than any one strain-cell line interaction but instead must be considered from an array of perspectives.

Another important question arising from our work is how ARHGEF26 localizes to the site of invasion during infection. Previous work speculated that ARHGEF26 binds phosphoinositides through its PH domain (7), however, our results do not support this model. Instead, we hypothesize that the previously described SCRIB-DLG1-ARHGEF26 complex (48) guides the GEF to the plasma membrane, possibly through DLG1’s ability to bind phosphoinositides (71). While phosphoinositide-binding plays a central role in both models, it is important to note that the former model was based on the premise that ARHGEF26 is involved in SopB-, but not SopE-mediated invasion. Instead, our data suggest that SopB and SopE cooperatively guide ARHGEF26 to the site of invasion, enabling ARHGEF26 to contribute to both SopB- and SopE-mediated invasion. How SopE contributes to this process is a mystery. It could be that SopE-mediated changes to phosphoinositides (9) are able to recruit the complex to the membrane, or some other positive feedback system may exist following SopE-mediated nucleotide exchange. Further, our data suggest that scaffolds other than DLG1 and SCRIB may be involved specifically in bringing active *ARHGEF26* to the SopB-induced membrane ruffle. Perhaps this DLG1/SCRIB-independent mechanism involves post-translational modifications to ARHGEF26, which have previously been reported to regulate ARHGEF26 activity (72).

The utilization of both a host and pathogen protein with similar functions (ARHGEF26 and SopE) to stimulate invasion is an unexpected and fascinating observation. Bacterial effectors that mimic host proteins often outperform their mammalian counterparts in order to promote pathogenesis (73, 74). That *S*. Typhi benefits from both the SopE GEF mimic and ARHGEF26 being present in HeLa cells is a striking exception to this paradigm, but not unheard of. Indeed, previous work has demonstrated that SopE acts cooperatively with the host GEF ARNO (CYTH2) to activate the WAVE complex and induce invasion (23). As we and others (19, 20) have provided evidence that RHOG may be dispensable for invasion, one potential reason for this cooperation may be differences in substrate specificity between SopE and ARHGEF26. Therefore, identifying the GTPase(s) that ARHGEF26 activates during infection represents an important future direction. One potential GTPase is RHOJ (also called TCL), a GTPase that has high homology to CDC42 (47) and is important *Salmonella* invasion (19). Little is known about RHOJ regulation, though it’s N-terminus has been shown to be critical for localization to the plasma membrane and for nucleotide exchange (75, 76). Future work will examine whether ARHGEF26 can facilitate nucleotide exchange with this or other GTPases.

Finally, we demonstrated that ARHGEF26 contributes to *S*. Typhimurium fitness in the enteric fever mouse model, as well as to *S*. Typhimurium-induced inflammation in the streptomycin pretreatment gastroenteritis model. This latter observation was surprising, as no report has shown *ARHGEF26* may promote proinflammatory processes, and in fact, some have speculated that it may be involved in anti-inflammatory processes. This is due to its relatively high levels in M2 macrophages (77) as well as its ability to suppress muramyl dipeptide-induced IL-8 production in NOD2 expressing HEK293 cells (78). While initially this latter finding appears to conflict with our results, we instead believe it merely reinforces a point implied by our data: ARHGEF26 has highly context dependent roles in regulating inflammation. For instance, ARHGEF26 is a strong regulator of IL-8 abundance in uninfected supernatant, but only a moderate regulator in *S*. Typhi and *S*. Typhimurium infected supernatant. The regulatory network is complicated further by our finding that ARHGEF26 likely regulates cytokine abundance through RHOG dependent and GEF independent mechanisms.

We speculate that there are two mechanisms by which ARHGEF26 could contribute to inflammation in the mouse gut. First, ARHGEF26 could impact proinflammatory cytokine release, as we observe in HeLa cells. Second, as ARHGEF26 has been shown to play a role in cell migration (72, 79), we speculate that *Arhgef26*^-/-^ mice may have reduced immune cell migration to the gut, leading to improved pathophysiology. Determining the role of ARHGEF26 during inflammation could have interesting implications on human heath, as inhibiting ARHGEF26 and/or RHOG could be a means of reducing inflammation-driven disease.

## Supporting information

SupplementalFile--MousePathologyScoring

## Acknowledgements and Funding

We would like to thank Alyson Barnes, Ben Schott, Alejandro Antonia, Sarah Jaslow, Kelly Pittman, Rachel Keener, and all other past and present members of the Ko Lab for their support throughout this project. In particular, we thank Kyle Gibbs for his thorough editing of this manuscript and frequent contributions to experimental design. We thank Dr. Keith Burridge for early discussion on ARHGEF26 and a gift of *Arhgef26^-/-^* mice. We also thank Dr. Stacy Horner and the Duke MGM Department for use of equipment. All schematic images were generated using Biorender.com. The *S*. Typhimurium *sopE*::*tet* strain was a gift from Heather Felise.

JSB was supported by National Institutes of Health 1F31AI143147. JSB, MIA, LW, and DCK were supported by National Institutes of Health R01AI118903 and R21AI144586. SA and RGM were supported by National Institutes of Health R01GM136826. The funders played no role in the study design, data collection and analysis, decision to publish, or preparation of the manuscript.

## Methods

Ethics Statement Work involving human lymphoblastoid cell lines has been reviewed by Duke Institutional Review Board and deemed to not constitute Human Subjects Research (Pro00044583, “Functional genetic screens of human variation using lymphoblastoid cell lines”). Mouse studies were carried out with approval by the Duke Institutional Animal Care and Use Committee (A145-18-06, “Analysis of genes affecting microbial virulence in mice”) and adhere to the *Guide for the Care and Use of Laboratory Animals* of the National Institutes of Health.

### Mammalian and Bacterial Cell Culture

HapMap LCLs (Coriell Institute) were cultured at 37°C in 5% CO_2_ in RPMI 1650 media (Invitrogen) supplemented with 10% FBS (Thermo-Fisher), 2 μM glutamine, 100 U/mL penicillin-G, and 100 mg/mL streptomycin. HeLa cells (Duke Cell Culture Facility) and Hek293T (Duke Cell Culture Facility) were grown in high glucose DMEM media supplemented with 10% FBS, 1mM glutamine, 100 U/mL penicillin-G, and 100mg/mL streptomycin. Cells used for *Salmonella* gentamicin protection assays were grown without antibiotics at least one hour prior to infection.

All *Salmonella* strains are derived from the *S*. Typhimurium strain 14028s or *S*. Typhi strain Ty2 and are listed in Table S2, and all plasmids are listed in Table S3. All knockout strains were generated by lambda red recombination (80). For infection of cells or mice, bacteria were grown overnight in LB broth (Miller formulation, BD), subcultured 1:33 in 1mL cultures, and grown for an additional two hours and forty minutes at 37°C shaking at 250 RPM. Strains with temperature sensitive plasmids were grown at 30°C and plasmids removed at 42°C. Ampicillin was added to LB at 100 μg/mL, kanamycin at 50 μg/mL.

### *Salmonella* Infection Assays

Infection assays were performed as previously described (17). Briefly, cells were infected with *S*. Typhimurium (LCLs MOI 30, 60 minutes infection. HeLa MOI 1, 30 minute infection, unless otherwise noted) or *S*. Typhi (LCLs MOI 10, 60 minute infection. HeLa MOI 30, 30 minute infection, unless otherwise noted). Post infection, cells were treated with 50 μg/mL gentamicin. Two hours post infection, IPTG was added to induce GFP expression. Three hours and fifteen minutes post infection, cells were stained with 7-aminoactinomycin D (Biomol) and analyzed on a Guava Easycyte Plus flow Cytometer (Millipore). Percent invasion was measured by quantifying the percent of GFP+ cells.

### Cellular GWAS Screen

Phenotypic screening in H2P2 on 528 LCLs and family-based GWAS analysis was performed using QFAM-parents with adaptive permutation in PLINK v1.9 (81) as previously described (17). All analyzed GWAS data is available through the H2P2 web atlas (http://h2p2.oit.duke.edu/H2P2Home/) (17). QQ plots were plotted using quantile-quantile function in R.

### Dual Luciferase Assay

The ARHGEF26 locus identified by H2P2 was cloned from the heterozygote HG02860 (population = Gambian and Western Divisions in the Gambia) into the pBV-Firefly Luciferase plasmid (39) by cut and paste cloning. The plasmid map is available here: https://benchling.com/s/seq-2427vsVPRqxsgOj5NM7P. Firefly luciferase plasmids and the Renilla luciferase plasmid pRL-SV40P (39) were co-transfected at a ratio of 50:1 into HeLa cells using the Lipofectamine 3000 kit (Thermo) according to manufacturer instructions. 48 hours post transfection, cells were lysed and analyzed for luciferase activity using the Dual-Luciferase Reporter Assay System (Promega). Luciferase activity was measured by a Synergy H1 plate reader (BioTek).

### siRNA Knockdown and Knockdown Confirmation

LCL (HG01697) knockdown was achieved by plating at 250,000 cells/well in a six well dish in 500 μL of Accell media (Dharmacon) with either non-targeting Accell siRNA #1 or an Accell *ARHGEF26* SMARTpool (1 μM total siRNA; Dharmacon). After 3 days, cells were resuspended in RPMI at 50,000 cells/well in a 96 well dish.

HeLa knockdown was performed using the following siRNA: siGenome Non-Targeting #5 or a siGENOME SMARTpool targeting *ARHGEF26, SCRIB, RHOG*, or *DLG1* (Horizon). siRNA were transfected into HeLa cells using the RNAi Max kit (Thermo) according to manufacturer instructions. Assays were performed forty-eight hours post infection as described above.

Simultaneously, knockdown was confirmed in each experiment by qPCR (Figure S1F). Briefly, RNA was harvested using a RNeasy kit (Qiagen), cDNA was generated with iScript (Bio-Rad), and qPCR was performed by using iTaq Universal Probes Supermix (Bio-Rad) and a QuantStudio 3 thermo cycler (Applied Biosystems). Primers are listed in Table S4. The cycling conditions were as follows: 95°C for 2 minutes, 95°C for 10 minutes, and 40 cycles of 95°C for 15 seconds followed by 60°C for 1 minute. All qPCR was run in technical duplicate or triplicate. The comparative threshold cycle (CT) was used to quantify transcripts, with the ribosomal 18s gene (RNA18S5) serving as the housekeeping control. ΔC_T_ values were calculated by subtracting the C_T_ value of the control gene from the target gene, and the ΔΔC_T_ was calculated by subtracting the nontargeting siRNA ΔC_T_ from the targeting siRNA ΔC_T_ value. Fold change represents 2^−ΔΔCT^.

### *ARHGEF26 and RHOG* Overexpression Plasmids

*ARHGEF26 and RHOG* overexpression plasmids (Table S3) were transformed using the Lipofectamine 3000 kit (Thermo) according to manufacturer instructions. Most plasmids were generated in previous work (45, 48), and all remaining plasmids were generated through site-directed mutagenesis (QuickChange Lightning, Agilent) or cut-and-paste cloning. Assays using overexpression plasmids were performed twenty-four hours post transfection.

### Microscopy

Cells were fixed for thirty minutes in 4% paraformaldehyde and blocked for thirty minutes in a 5% normal donkey serum, 0.2% saponin, PBS solution. Cells were incubated overnight at 4°C with an anti-myc antibody (Developmental Studies Hybridoma Bank, 9e10, followed by secondary staining using Alexa Fluor™ secondary anti-mouse antibody (Thermo). Anti-Myc (9e10) was deposited to the DSHB by Bishop, J.M. (DSHB Hybridoma Product 9e10). Actin staining was performed using Alexa Fluor™ 647 Phalloidin (Thermo) according to manufacturer instructions. Micrographs were taken using an AMG EVOS microscope.

### Phosphoinositide Dot Blot Assays

Twenty-four hours before transfection, 1,500,000 HeLa cells were plated on a 10-cm dish or 1,000,000 Hek293T cells were plated in three separate wells of a six-well dish. Cells were transfected as above, but to normalize for expression, 1μg of AKT-PH-GFP was diluted with 7μg vector. Twenty-four hours later, cells were washed, and directly scraped into lysis buffer (50mM Tris, pH 7.6, 150mM NaCl, 1% Triton X-100, 5mM MgCl2, cOmplete Mini protease inhibitor cocktail (Sigma)), and incubated at 4°C for 30 minutes. PIP strips (PIP Strips (Echelon) were blocked using Odyssey^®^ Blocking Buffer (Licor) or Intercept Blocking Buffer (Licor). Samples were cleared by centrifugation and diluted 1:25 into blocking buffer before addition to the PIP strips (Echelon Biosciences). After one hour incubation with rocking at room temperature, PIP strips were washed 3 times with PBS-T, and incubated with an anti-GFP primary antibody (Novus, NB600-308). After one hour rocking at room temperature, PIP strips were washed three times with PBS-T, and stained with a IRDye^®^ donkey anti-rabbit secondary antibody (Licor). After thirty minutes, strips were washed three times with PBS-T, once with PBS, and imaged on a LI-COR Odyssey Classic.

### Analysis of HeLa Cytokine Production

For siRNA experiments, two days post transfection media was changed 2 hours before infection. HeLa cells were then infected with late log phase bacteria (*S*. Typhimurium: MOI 1; *S*. Typhi: MOI 30) for 60 minutes. After 60 minutes, gentamycin was added and bacteria were returned to 37°C incubator for 5 hours. Six hours post infection supernatants were collected. For overexpression experiments, media was changed 18-24 hours post infection. Six hours after the media change, supernatants were collected. Supernatants were stored at −80°C until use. Cytokine concentrations were determined using a human IL-8 DuoSet ELISA kit (R&D Systems).

### Mouse Infections

C57BL6/J mice were obtained from JAX and housed in barrier cages in the Duke University’s Division of Laboratory Animal Resources husbandry facility. Following arrival at the Duke University’s Division of Laboratory Animal Resources husbandry facility, *Arhgef26^-/-^* mice (64) were rederived as specific pathogen free mice by embryo transplantation. Mice were fed rodent diet 5053 chow.

For the enteric fever model of infection, age and sex matched 7-12 week old C57BL/6J or *Arhgef26^-/-^* mice were fasted for 12 hours prior to infection, and treated with a 100μL of a 10% sodium bicarbonate solution 30 minutes prior to infection. Bacteria were grown as described above, washed, and resuspended in PBS at a concentration of 1×10^9^ bacteria/mL, and 100μL were administered to the mice for an estimated final dose of 1×10^8^ bacteria/mouse. Inoculum was confirmed by plating for CFUs. All mice were monitored daily for changes in morbidity. Mice were euthanized by CO_2_ asphyxiation four days post infection and tissues were harvested, weighed, homogenized, and plated for CFU quantification.

For the gastroenteritis model of infection (10), mice were 7-12 week old C57BL/6J or *Arhgef26^-/-^* mice were fasted four hours before treatment with 20μg of streptomycin (Sigma) in 75μL of sterile water 24 hours before infection. Food was returned until four hours before infection, when they were fasted again. Thirty minutes before infection, mice received 100μL of a 10% sodium bicarbonate solution. Bacteria, grown as described above, were washed and resuspended in PBS at a concentration of 1×10^10^ bacteria/mL, and 100μL were administered to the mice for an estimated final dose of 1×10^9^ bacteria/mouse. Food was returned four hours after infection. Inoculum was confirmed by plating for CFUs. All mice were monitored daily for changes in morbidity. Two days post infection, mice were euthanized by CO_2_ asphyxiation and tissues were removed either weighed and plated for CFUs as described above or prepared for histopathologic examination.

Cecal and colon tissues were fixed 48-72 hours in 10% neutral buffered formalin, processed routinely, embedded in paraffin, cut at 5mm and stained with hematoxylin and eosin. Tissues were evaluated in a masked fashion by a board-certified veterinary pathologist (JIE) with allocation group concealment. Tissues were scored using a semi-quantitative grading system of multiple parameters and anatomic compartments (lumen, surface epithelium, mucosa, and submucosa) to assign summary pathologic injury scores (82).

The histopathologic scoring was (scores in parenthesis). (a) Lumen: empty (0), necrotic epithelial cells (scant, 1; moderate, 2; dense, 3), and polymorphonuclear leukocytes (PMNs) (scant, 2; moderate, 3; dense, 4). (b) Surface epithelium: No pathological changes (0); mild, moderate, or severe regenerative changes (1, 2, or 3, respectively); patchy or diffuse desquamation (1 or 2); PMNs in epithelium (1); and ulceration (1). (c) Mucosa: No pathological changes (0); rare (<15%), moderate (15 to 50%), or abundant (>50%) crypt abscesses (1, 2, or 3, respectively); presence of mucinous plugs (1); presence of granulation tissue (1). (d) Submucosa: No pathological changes (0); mononuclear cell infiltrate (1 small aggregate, <1 aggregate, or large aggregates plus increased single cells) (0, 1, or 2, respectively); PMN infiltrate (no extravascular PMNs, single extravascular PMNs, or PMN aggregates) (0, 1, or 2, respectively); mild, moderate, or severe edema (0, 1, or 2, respectively).

### Statistics

All statistics were performed using Graphpad Prism 8 or Microsoft Excel, unless otherwise noted. Bars and central tendencies representations as well as p-value calculations are described in all figure legends. Where noted, inter-experimental noise was removed prior to data visualization or statistical analysis by standardizing data to the grand mean by multiplying values within an experiment by a constant (average of all experiments divided by average of specific experiment). Data points were only excluded if technical failure could be proven (*i.e*. failed RNAi knockdown measured by qPCR), or if identified by an outlier test. If datapoints were removed by an outlier test, the original results are reported in the figure legend.

### Data Availability

All cellular GWAS data from H2P2 are available through the H2P2 web atlas (http://h2p2.oit.duke.edu/H2P2Home/) (17). All other relevant data are within the manuscript and its Supporting Information files. All plasmids, primers, bacterial strains, and mice are available upon request. No other tools or datasets were generated in this manuscript.

## References

1. Lewis MR. The Ormation of Vacuoles Due to Bacillus Typhosus in the Cells of Tissue Cultures of the Intestine of the Chick Embryo. J Exp Med. 1920;31(3):293–311.

2. Salmonella interactions with host cells: type III secretion at work.,(2001).

3. Cloning and molecular characterization of genes whose products allow Salmonella typhimurium to penetrate tissue culture cells., (1989).

4. Hardt WD, Chen LM, Schuebel KE, Bustelo XR, Galan JE. S. typhimurium encodes an activator of Rho GTPases that induces membrane ruffling and nuclear responses in host cells. Cell. 1998;93(5):815–26.

5. Hernandez LD, Hueffer K, Wenk MR, Galan JE. Salmonella modulates vesicular traffic by altering phosphoinositide metabolism. Science. 2004;304(5678):1805–7.

6. Mallo GV, Espina M, Smith AC, Terebiznik MR, Aleman A, Finlay BB, et al. SopB promotes phosphatidylinositol 3-phosphate formation on Salmonella vacuoles by recruiting Rab5 and Vps34. J Cell Biol. 2008;182(4):741–52.

7. Patel JC, Galan JE. Differential activation and function of Rho GTPases during Salmonella-host cell interactions. J Cell Biol. 2006;175(3):453–63.

8. Terebiznik MR, Vieira OV, Marcus SL, Slade A, Yip CM, Trimble WS, et al. Elimination of host cell PtdIns(4,5)P(2) by bacterial SigD promotes membrane fission during invasion by Salmonella. Nat Cell Biol. 2002;4(10):766–73.

9. Zhou D, Chen LM, Hernandez L, Shears SB, Galan JE. A Salmonella inositol polyphosphatase acts in conjunction with other bacterial effectors to promote host cell actin cytoskeleton rearrangements and bacterial internalization. Mol Microbiol. 2001;39(2):248–59.

10. Hapfelmeier S, Ehrbar K, Stecher B, Barthel M, Kremer M, Hardt WD. Role of the Salmonella pathogenicity island 1 effector proteins SipA, SopB, SopE, and SopE2 in Salmonella enterica subspecies 1 serovar Typhimurium colitis in streptomycin-pretreated mice. Infect Immun. 2004;72(2):795–809.

11. Pretreatment of mice with streptomycin provides a Salmonella enterica serovar Typhimurium colitis model that allows analysis of both pathogen and host., (2003).

12. Jin C, Gibani MM, Moore M, Juel HB, Jones E, Meiring J, et al. Efficacy and immunogenicity of a Vi-tetanus toxoid conjugate vaccine in the prevention of typhoid fever using a controlled human infection model of Salmonella Typhi: a randomised controlled, phase 2b trial. Lancet. 2017;390(10111):2472–80.

13. Dunstan SJ, Hue NT, Han B, Li Z, Tram TT, Sim KS, et al. Variation at HLA-DRB1 is associated with resistance to enteric fever. Nat Genet. 2014;46(12):1333–6.

14. Gilchrist JJ, Rautanen A, Fairfax BP, Mills TC, Naranbhai V, Trochet H, et al. Risk of nontyphoidal Salmonella bacteraemia in African children is modified by STAT4. Nat Commun. 2018;9(1):1014.

15. Ko DC, Gamazon ER, Shukla KP, Pfuetzner RA, Whittington D, Holden TD, et al. Functional genetic screen of human diversity reveals that a methionine salvage enzyme regulates inflammatory cell death. Proc Natl Acad Sci U S A. 2012;109(35):E2343–52.

16. Ko DC, Shukla KP, Fong C, Wasnick M, Brittnacher MJ, Wurfel MM, et al. A genome-wide in vitro bacterial-infection screen reveals human variation in the host response associated with inflammatory disease. Am J Hum Genet. 2009;85(2):214–27.

17. Wang L, Pittman KJ, Barker JR, Salinas RE, Stanaway IB, Williams GD, et al. An Atlas of Genetic Variation Linking Pathogen-Induced Cellular Traits to Human Disease. Cell Host Microbe. 2018;24(2):308–23 e6.

18. Alvarez MI, Glover LC, Luo P, Wang L, Theusch E, Oehlers SH, et al. Human genetic variation in VAC14 regulates Salmonella invasion and typhoid fever through modulation of cholesterol. Proc Natl Acad Sci U S A. 2017;114(37):E7746–E55.

19. Truong D, Boddy KC, Canadien V, Brabant D, Fairn GD, D’Costa VM, et al. Salmonella exploits host Rho GTPase signalling pathways through the phosphatase activity of SopB. Cell Microbiol. 2018;20(10):e12938.

20. Aiastui A, Pucciarelli MG, Garcia-del Portillo F. Salmonella enterica serovar typhimurium invades fibroblasts by multiple routes differing from the entry into epithelial cells. Infect Immun. 2010;78(6):2700–13.

21. Hume PJ, Singh V, Davidson AC, Koronakis V. Swiss Army Pathogen: The Salmonella Entry Toolkit. Front Cell Infect Microbiol. 2017;7:348.

22. Patel JC, Galan JE. Manipulation of the host actin cytoskeleton by Salmonella--all in the name of entry. Curr Opin Microbiol. 2005;8(1):10–5.

23. Humphreys D, Davidson A, Hume PJ, Koronakis V. Salmonella virulence effector SopE and Host GEF ARNO cooperate to recruit and activate WAVE to trigger bacterial invasion. Cell Host Microbe. 2012;11(2):129–39.

24. Humphreys D, Davidson AC, Hume PJ, Makin LE, Koronakis V. Arf6 coordinates actin assembly through the WAVE complex, a mechanism usurped by Salmonella to invade host cells. Proc Natl Acad Sci U S A. 2013;110(42):16880–5.

25. Patel JC, Galan JE. Investigating the function of Rho family GTPases during Salmonella/host cell interactions. Methods Enzymol. 2008;439:145–58.

26. Chen LM, Hobbie S, Galan JE. Requirement of CDC42 for Salmonella-induced cytoskeletal and nuclear responses. Science. 1996;274(5295):2115–8.

27. Stender S, Friebel A, Linder S, Rohde M, Mirold S, Hardt WD. Identification of SopE2 from Salmonella typhimurium, a conserved guanine nucleotide exchange factor for Cdc42 of the host cell. Mol Microbiol. 2000;36(6):1206–21.

28. Unsworth KE, Way M, McNiven M, Machesky L, Holden DW. Analysis of the mechanisms of Salmonella-induced actin assembly during invasion of host cells and intracellular replication. Cell Microbiol. 2004;6(11):1041–55.

29. Ablain J, Xu M, Rothschild H, Jordan RC, Mito JK, Daniels BH, et al. Human tumor genomics and zebrafish modeling identify SPRED1 loss as a driver of mucosal melanoma. Science. 2018;362(6418):1055–60.

30. Lilic M, Galkin VE, Orlova A, VanLoock MS, Egelman EH, Stebbins CE. Salmonella SipA polymerizes actin by stapling filaments with nonglobular protein arms. Science. 2003;301(5641):1918–21.

31. Hayward RD, Koronakis V. Direct nucleation and bundling of actin by the SipC protein of invasive Salmonella. EMBO J. 1999;18(18):4926–34.

32. Zhou D, Mooseker MS, Galan JE. An invasion-associated Salmonella protein modulates the actin-bundling activity of plastin. Proc Natl Acad Sci U S A. 1999;96(18):10176–81.

33. Francis CL, Starnbach MN, Falkow S. Morphological and cytoskeletal changes in epithelial cells occur immediately upon interaction with Salmonella typhimurium grown under low-oxygen conditions. Mol Microbiol. 1992;6(21):3077–87.

34. Criss AK, Casanova JE. Coordinate regulation of Salmonella enterica serovar Typhimurium invasion of epithelial cells by the Arp2/3 complex and Rho GTPases. Infect Immun. 2003;71(5):2885–91.

35. Machiela MJ, Chanock SJ. LDlink: a web-based application for exploring population-specific haplotype structure and linking correlated alleles of possible functional variants. Bioinformatics. 2015;31(21):3555–7.

36. Ward LD, Kellis M. HaploReg v4: systematic mining of putative causal variants, cell types, regulators and target genes for human complex traits and disease. Nucleic Acids Res. 2016;44(D1):D877–81.

37. Consortium GT. The Genotype-Tissue Expression (GTEx) project. Nat Genet. 2013;45(6):580–5.

38. Lappalainen T, Sammeth M, Friedlander MR, t Hoen PA, Monlong J, Rivas MA, et al. Transcriptome and genome sequencing uncovers functional variation in humans. Nature. 2013;501(7468):506–11.

39. He TC, Chan TA, Vogelstein B, Kinzler KW. PPARdelta is an APC-regulated target of nonsteroidal anti-inflammatory drugs. Cell. 1999;99(3):335–45.

40. Hanisch J, Ehinger J, Ladwein M, Rohde M, Derivery E, Bosse T, et al. Molecular dissection of Salmonella-induced membrane ruffling versus invasion. Cell Microbiol. 2010;12(1):84–98.

41. Pruim RJ, Welch RP, Sanna S, Teslovich TM, Chines PS, Gliedt TP, et al. LocusZoom: regional visualization of genome-wide association scan results. Bioinformatics. 2010;26(18):2336–7.

42. Boulter E, Garcia-Mata R, Guilluy C, Dubash A, Rossi G, Brennwald PJ, et al. Regulation of Rho GTPase crosstalk, degradation and activity by RhoGDI1. Nat Cell Biol. 2010;12(5):477–83.

43. Ren XD, Kiosses WB, Schwartz MA. Regulation of the small GTP-binding protein Rho by cell adhesion and the cytoskeleton. EMBO J. 1999;18(3):578–85.

44. Garcia-Mata R, Boulter E, Burridge K. The ‘invisible hand’: regulation of RHO GTPases by RHOGDIs. Nature reviews Molecular cell biology. 2011;12(8):493–504.

45. Ellerbroek SM, Wennerberg K, Arthur WT, Dunty JM, Bowman DR, DeMali KA, et al. SGEF, a RhoG guanine nucleotide exchange factor that stimulates macropinocytosis. Mol Biol Cell. 2004;15(7):3309–19.

46. Reinhard NR, Van Der Niet S, Chertkova A, Postma M, Hordijk PL, Gadella TWJ, Jr., et al. Identification of guanine nucleotide exchange factors that increase Cdc42 activity in primary human endothelial cells. Small GTPases. 2019:1–15.

47. Vignal E, De Toledo M, Comunale F, Ladopoulou A, Gauthier-Rouviere C, Blangy A, et al. Characterization of TCL, a new GTPase of the rho family related to TC10 andCcdc42. J Biol Chem. 2000;275(46):36457–64.

48. Awadia S, Huq F, Arnold TR, Goicoechea SM, Sun YJ, Hou T, et al. SGEF forms a complex with Scribble and Dlg1 and regulates epithelial junctions and contractility. J Cell Biol. 2019;218(8):2699–725.

49. Krishna Subbaiah V, Massimi P, Boon SS, Myers MP, Sharek L, Garcia-Mata R, et al. The invasive capacity of HPV transformed cells requires the hDlg-dependent enhancement of SGEF/RhoG activity. PLoS Pathog. 2012;8(2):e1002543.

50. Lemmon MA. Pleckstrin homology (PH) domains and phosphoinositides. Biochem Soc Symp. 2007(74):81–93.

51. Marcus SL, Wenk MR, Steele-Mortimer O, Finlay BB. A synaptojanin-homologous region of Salmonella typhimurium SigD is essential for inositol phosphatase activity and Akt activation. FEBS letters. 2001;494(3):201–7.

52. Mason D, Mallo GV, Terebiznik MR, Payrastre B, Finlay BB, Brumell JH, et al. Alteration of epithelial structure and function associated with PtdIns(4,5)P2 degradation by a bacterial phosphatase. J Gen Physiol. 2007;129(4):267–83.

53. Norris FA, Wilson MP, Wallis TS, Galyov EE, Majerus PW. SopB, a protein required for virulence of Salmonella dublin, is an inositol phosphate phosphatase. Proc Natl Acad Sci U S A. 1998;95(24):14057–9.

54. Yu JW, Mendrola JM, Audhya A, Singh S, Keleti D, DeWald DB, et al. Genome-wide analysis of membrane targeting by S. cerevisiae pleckstrin homology domains. Molecular cell. 2004;13(5):677–88.

55. Skowronek KR, Guo F, Zheng Y, Nassar N. The C-terminal basic tail of RhoG assists the guanine nucleotide exchange factor trio in binding to phospholipids. J Biol Chem. 2004;279(36):37895–907.

56. Narayan K, Lemmon MA. Determining selectivity of phosphoinositide-binding domains. Methods. 2006;39(2):122–33.

57. Gewirtz AT, Rao AS, Simon PO, Jr., Merlin D, Carnes D, Madara JL, et al. Salmonella typhimurium induces epithelial IL-8 expression via Ca(2+)-mediated activation of the NF-kappaB pathway. The Journal of clinical investigation. 2000;105(1):79–92.

58. Huang FC, Werne A, Li Q, Galyov EE, Walker WA, Cherayil BJ. Cooperative interactions between flagellin and SopE2 in the epithelial interleukin-8 response to Salmonella enterica serovar typhimurium infection. Infect Immun. 2004;72(9):5052–62.

59. Bruno VM, Hannemann S, Lara-Tejero M, Flavell RA, Kleinstein SH, Galan JE. Salmonella Typhimurium type III secretion effectors stimulate innate immune responses in cultured epithelial cells. PLoS Pathog. 2009;5(8):e1000538.

60. Hobbie S, Chen LM, Davis RJ, Galan JE. Involvement of mitogen-activated protein kinase pathways in the nuclear responses and cytokine production induced by Salmonella typhimurium in cultured intestinal epithelial cells. J Immunol. 1997;159(11):5550–9.

61. Sun H, Kamanova J, Lara-Tejero M, Galan JE. Salmonella stimulates pro-inflammatory signalling through p21-activated kinases bypassing innate immune receptors. Nat Microbiol. 2018;3(10):1122–30.

62. Keestra AM, Winter MG, Auburger JJ, Frassle SP, Xavier MN, Winter SE, et al. Manipulation of small Rho GTPases is a pathogen-induced process detected by NOD1. Nature. 2013;496(7444):233–7.

63. Hobert ME, Sands KA, Mrsny RJ, Madara JL. Cdc42 and Rac1 regulate late events in Salmonella typhimurium-induced interleukin-8 secretion from polarized epithelial cells. J Biol Chem. 2002;277(52):51025–32.

64. Samson T, van Buul JD, Kroon J, Welch C, Bakker EN, Matlung HL, et al. The guanine-nucleotide exchange factor SGEF plays a crucial role in the formation of atherosclerosis. PLoS One. 2013;8(1):e55202.

65. Yeung ATY, Choi YH, Lee AHY, Hale C, Ponstingl H, Pickard D, et al. A Genome-Wide Knockout Screen in Human Macrophages Identified Host Factors Modulating Salmonella Infection. mBio. 2019;10(5).

66. Van Puyvelde S, Pickard D, Vandelannoote K, Heinz E, Barbe B, de Block T, et al. An African Salmonella Typhimurium ST313 sublineage with extensive drug-resistance and signatures of host adaptation. Nat Commun. 2019;10(1):4280.

67. Jaslow SL, Gibbs KD, Fricke WF, Wang L, Pittman KJ, Mammel MK, et al. Salmonella Activation of STAT3 Signaling by SarA Effector Promotes Intracellular Replication and Production of IL-10. Cell Rep. 2018;23(12):3525–36.

68. Misselwitz B, Barrett N, Kreibich S, Vonaesch P, Andritschke D, Rout S, et al. Near surface swimming of Salmonella Typhimurium explains target-site selection and cooperative invasion. PLoS Pathog. 2012;8(7):e1002810.

69. Zhang K, Riba A, Nietschke M, Torow N, Repnik U, Putz A, et al. Minimal SPI1-T3SS effector requirement for Salmonella enterocyte invasion and intracellular proliferation in vivo. PLoS Pathog. 2018;14(3):e1006925.

70. Fattinger SA, Bock D, Di Martino ML, Deuring S, Samperio Ventayol P, Ek V, et al. Salmonella Typhimurium discreet-invasion of the murine gut absorptive epithelium. PLoS Pathog. 2020;16(5):e1008503.

71. Awad A, Sar S, Barre R, Cariven C, Marin M, Salles JP, et al. SHIP2 regulates epithelial cell polarity through its lipid product, which binds to Dlg1, a pathway subverted by hepatitis C virus core protein. Mol Biol Cell. 2013;24(14):2171–85.

72. Okuyama Y, Umeda K, Negishi M, Katoh H. Tyrosine Phosphorylation of SGEF Regulates RhoG Activity and Cell Migration. PLoS One. 2016;11(7):e0159617.

73. LaRock DL, Brzovic PS, Levin I, Blanc MP, Miller SI. A Salmonella typhimurium-translocated glycerophospholipid:cholesterol acyltransferase promotes virulence by binding to the RhoA protein switch regions. J Biol Chem. 2012;287(35):29654–63.

74. Gibbs KD, Washington EJ, Jaslow SL, Bourgeois JS, Foster MW, Guo R, et al. The Salmonella Secreted Effector SarA/SteE Mimics Cytokine Receptor Signaling to Activate STAT3. Cell Host Microbe. 2020;27(1):129–39 e4.

75. Ackermann KL, Florke RR, Reyes SS, Tader BR, Hamann MJ. TCL/RhoJ Plasma Membrane Localization and Nucleotide Exchange Is Coordinately Regulated by Amino Acids within the N Terminus and a Distal Loop Region. J Biol Chem. 2016;291(45):23604–17.

76. Florke RR, Young GT, Hamann MJ. Unraveling a model of TCL/RhoJ allosterism using TC10 reverse chimeras. Small GTPases. 2020;11(2):138–45.

77. Redka DS, Gutschow M, Grinstein S, Canton J. Differential ability of proinflammatory and anti-inflammatory macrophages to perform macropinocytosis. Mol Biol Cell. 2018;29(1):53–65.

78. Warner N, Burberry A, Pliakas M, McDonald C, Nunez G. A genome-wide small interfering RNA (siRNA) screen reveals nuclear factor-kappaB (NF-kappaB)-independent regulators of NOD2-induced interleukin-8 (IL-8) secretion. J Biol Chem. 2014;289(41):28213–24.

79. Valdivia A, Goicoechea SM, Awadia S, Zinn A, Garcia-Mata R. Regulation of circular dorsal ruffles, macropinocytosis, and cell migration by RhoG and its exchange factor, Trio. Mol Biol Cell. 2017;28(13):1768–81.

80. Datsenko KA, Wanner BL. One-step inactivation of chromosomal genes in Escherichia coli K-12 using PCR products. Proc Natl Acad Sci U S A. 2000;97(12):6640–5.

81. Chang CC, Chow CC, Tellier LC, Vattikuti S, Purcell SM, Lee JJ. Second-generation PLINK: rising to the challenge of larger and richer datasets. GigaScience. 2015;4:7.

82. Coburn B, Li Y, Owen D, Vallance BA, Finlay BB. Salmonella enterica serovar Typhimurium pathogenicity island 2 is necessary for complete virulence in a mouse model of infectious enterocolitis. Infect Immun. 2005;73(6):3219–27.

