## SupplementalFiguresandTables for "ARHGEF26 enhances *Salmonella* invasion and inflammation in cells and mice"

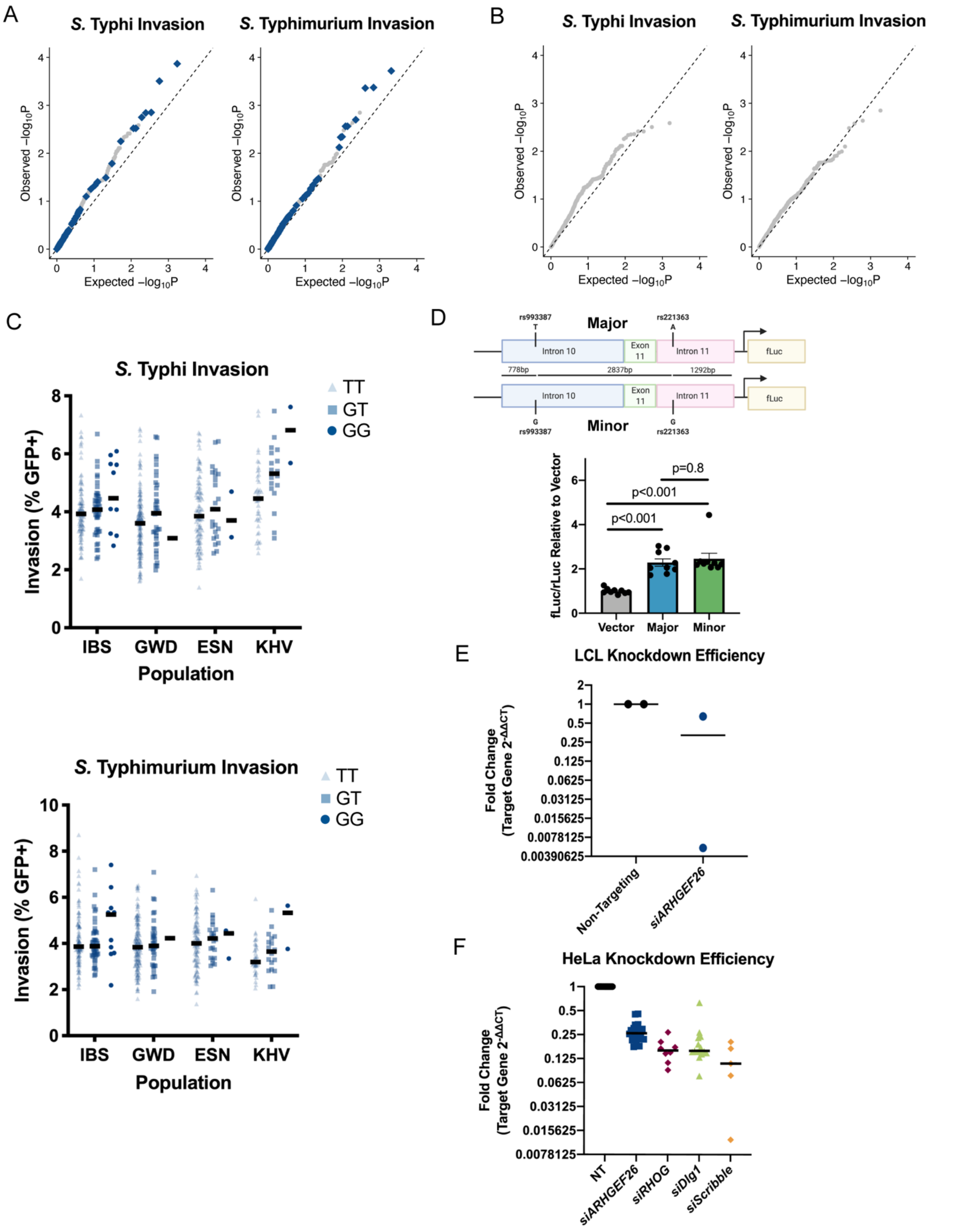

**Figure S1: Natural variation in *ARHGEF26* and *ARHGEF26* knockdown associate with reduced *Salmonella* invasion**. (A) Stratified QQ plots of invasion reveal that, in addition to rs993387, multiple SNPs in *ARHGEF26*

(blue diamonds) associate with *S.* Typhi or *S.* Typhimurium invasion in H2P2 at lower p-values than expected by chance. (B) Removing SNPs in *ARHGEF26* from the QQ plot removes any deviation of SNPs in SPI-1 associated genes from p-values expected by chance. (C) rs993387 association was observed across all four populations studied in H2P2 (IBS, Iberians from Spain; GWD, Gambian from the Western Divisions of The Gambia; ESN, Esan in Nigeria; KHV, Kinh in Ho Chi Minh City, Vietnam). Each dot represents a single LCL line averaged across three independent experiments. Black bar represents the median. LCLs in each population follow the trend of TT < GT < GG, except in GWD and ESN for Typhi where only 1 or 2 GG individuals were assayed. (E) A luciferase reporter system was generated to assess whether the rs993387 locus has enhancer activity. A roughly 5kb region was cloned from a heterozygous individual (HG02860, GWD) into pBV-Luc upstream of a minimal promoter and the firefly luciferase gene. Performing a dual luciferase experiment in HeLa cells revealed enhanced luciferase expression with the rs993387 locus. Bars represent the relative firefly luciferase/renilla luciferase activity, with vector set to 1. P-Values generated from one-way ANOVA with Tukey's multiple comparisons test on the log transformed values. (F) siRNA targeting *ARHGEF26* results in reduced expression in LCLs (HG01697, IBS). (G) siRNA targeting *ARHGEF26, DLG1, SCRIB,* and *RHOG* results in reduced expression in HeLa cells. Lines in F and G represent median fold change. Fold change is calculated as $2^{-\Delta\Delta CT}$ using RNA18S5 as a housekeeping control gene.

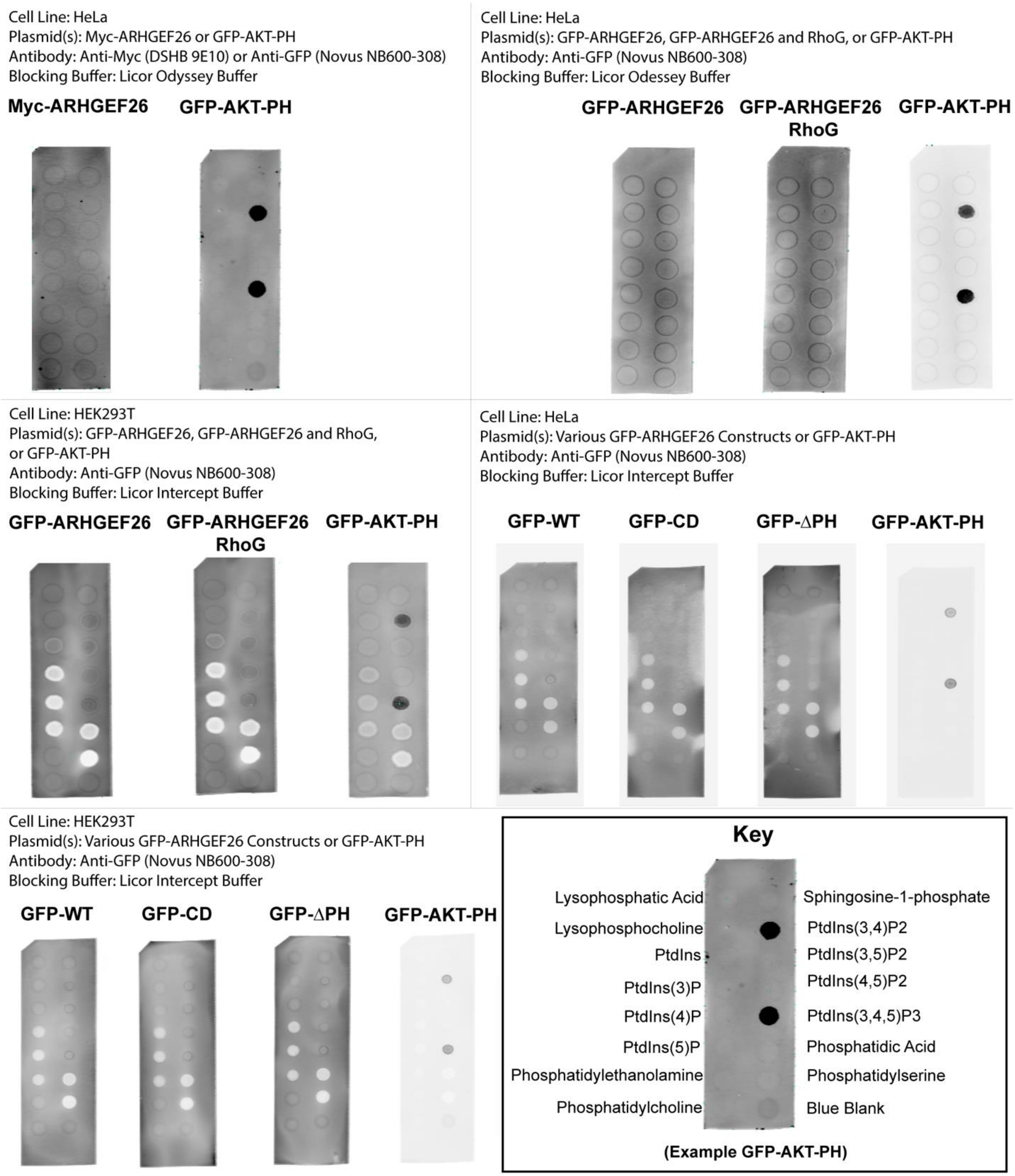

**Figure S2: ARHGEF26 does not demonstrate phosphoinositide binding.** ARHGEF26, RHOG, and the GFP-AKT-PH constructs were overexpressed in either HeLa or Hek293T cells. Cells were lysed, and protein extract was diluted in the listed blocking buffer before incubation on PIP strips. No robust ARHGEF26 signal on dotted phosphoinositide species could be detected following immunostaining. Key in bottom right displays location of each phosphoinositide species using a PIP strip incubated with GFP-AKT-PH (also shown in top left) as an example.

**Supplemental Table 1: Genes used for stratification.**

| Gene | Complex | Justification |
|---|---|---|
| *ABI1* | WAVE | (1-3) |
| *ABI2* | WAVE | (1-3) |
| *ABI3* | WAVE | (1-3) |
| *ACTB* | | (4-7) |
| *ACTR2* | ARP2/3 | (1, 2, 8) |
| *ACTR3* | ARP2/3 | (1, 2, 8) |
| *ARF1* | | (3) |
| *ARF6* | | (3, 9) |
| *ARHGEF26* (*SGEF*) | | (10) |
| *ARPC1A* | ARP2/3 | (1, 2, 8) |
| *ARPC1B* | ARP2/3 | (1, 2, 8) |
| *ARPC2* | ARP2/3 | (1, 2, 8) |
| *ARPC3* | ARP2/3 | (1, 2, 8) |
| *ARPC4* | ARP2/3 | (1, 2, 8) |
| *ARPC5* | ARP2/3 | (1, 2, 8) |
| *BRK1* | WAVE | (1-3) |
| *CDC42* | | (2, 8, 10, 11) |
| *CYFIP1* | WAVE | (1-3) |
| *CYFIP2* | WAVE | (1-3) |
| *CYTH2* (*ARNO*) | | (3, 9) |
| *NCKAP1* | WAVE | (1-3) |
| *RAC1* | | (2, 10-12) |
| *RHOG* | | (10, 13) |
| *WASF1* | WAVE | (1-3) |
| *WASF2* | WAVE | (1-3) |

**Supplemental Table 2: Bacterial strains used in this study**

| Designation | Serovar | Genotype | Plasmid | Resistance |
|---|---|---|---|---|
| CS093 | *S.* Typhimurium | Wild-Type (14028s) | | |
| DCK22 | *S.* Typhimurium | Wild-Type | p67GFP3.1 | Ampicillin |
| DCK483 | *S.* Typhimurium | Wild-Type | pWSK29 | Ampicillin |
| DCK484 | *S.* Typhimurium | Wild-Type | pWSK129 | Kanamycin |

| | | | | |
|---|---|---|---|---|
| DCK89 | *S.* Typhimurium | Δ*SopE2::tetR* | | Tetracycline |
| DCK95 | *S.* Typhimurium | Δ*SopE2::tetR* | p67GFP3.1 | Tetracycline, Ampicillin |
| DCK103 | *S.* Typhimurium | Δ*sopB* | | |
| DCK768 | *S.* Typhimurium | Δ*sopB* | pWSK29 | Ampicillin |
| DCK783 | *S.* Typhimurium | Δ*sopB* | pWSK129 | Kanamycin |
| DCK971 | *S.* Typhimurium | Δ*prgH* | pWSK129 | Kanamycin |
| CS092 | *S.* Typhi | Wild-Type (Ty2) | | |
| DCK33 | *S.* Typhi | Wild-Type (Ty2) | p67GFP3.1 | Ampicillin |
| DCK305 | *S.* Typhi | Δ*sopB* | | |
| DCK308 | *S.* Typhi | Δ*sopB* | p67GFP3.1 | Ampicillin |
| DCK306 | *S.* Typhi | ΔsopE | | |
| DCK309 | *S.* Typhi | ΔsopE | p67GFP3.1 | Ampicillin |
| DCK307 | *S.* Typhi | Δ*prgH* | | |
| DCK310 | *S.* Typhi | Δ*prgH* | p67GFP3.1 | Ampicillin |

| **Supplemental Table 3: Plasmids used in this study** | | | | |
|---|---|---|---|---|
| **Bacterial Stock** | **Parental Plasmid** | **Insert** | **Resistance** | **Source** |
| DCK18 | p67GFP3.1 | | Ampicillin | (14) |
| DCK482 | pWSK29 | | Ampicillin | (15) |
| DCK827 | pWSK129 | | Kanamycin | (15) |
| CS943 | pCP20 | | Ampicillin | (16) |
| CS946 | pKD4 | | Kanamycin | (16) |
| HB2502 | pKD46 | | Ampicillin | (16) |
| DCK718 | pCMV-Myc | | Ampicillin | (17) |
| DCK719 | pCMV-Myc | ARHGEF26 | Ampicillin | (17) |
| DCK749 | pCMV-Myc | ARHGEF26 ΔETNV (Δaa868-871) | Ampicillin | (18) |
| DCK750 | pCMV-Myc | ARHGEF26 aa1-400 | Ampicillin | (18) |
| DCK751 | pCMV-Myc | ARHGEF26 Catalytiacally Dead (R446A, N621A) | Ampicillin | (18) |
| DCK752 | pCMV-Myc | ARHGEF26 aa414-871 | Ampicillin | (18) |
| DCK753 | pCMV-Myc | ARHGEF26 PH-DH domain (aa431-792) | Ampicillin | (18) |
| DCK754 | pCMV-Myc | ARHGEF26 ΔSH3 (Δaa787-871) | Ampicillin | (18) |
| DCK757 | pCMV-Myc | ARHGEF26 ΔPH (Δaa656-726) | Ampicillin | This Study |
| DCK53 | pEGFP-C1 | | Kanamycin | Clontech |
| DCK835 | pEGFP-C1 | FLAG-ARHGEF26 | Kanamycin | This Study |

| | | | | |
|---|---|---|---|---|
| DCK837 | pEGFP-C1 | FLAG-ARHGEF26 Catalytiacally Dead (R446A, N621A) | Kanamycin | This Study |
| DCK839 | pEGFP-C1 | FLAG-ARHGEF26 ∆PH (∆∆aa656-726) | Kanamycin | This Study |
| DCK841 | pEGFP-C1 | RHOG | Kanamycin | This Study |
| DCK843 | pEGFP-C1 | RHOG Q61L (Constitutively Active) | Kanamycin | This Study |
| DCK784 | pCS2 | RHOG | Ampicillin | This Study |
| DCK531 | pBVLuc | | Ampicillin | (19) |
| DCK1013 | pBVLuc | ARHGEF26 rs993387 Locus Major Allele (HG02860) | Ampicillin | This Study |
| DCK1014 | pBVLuc | ARHGEF26 rs993387 Locus Minor Allele (HG02860) | Ampicillin | This Study |
| DCK534 | pRL-SV40P | | Ampicillin | (19) |

| **Supplemental Table 4: Oligonucleotides** | | | |
|---|---|---|---|
| ***Taqman Assays*** | | | |
| **Target Gene** | **Assay ID** | **Source** | |
| *ARHGEF26* | Hs00248943_m1 | ThermoFisher | |
| *SCRIB* | Hs00363005_m1 | ThermoFisher | |
| *DLG1* | Hs00938204_m1 | ThermoFisher | |
| *RHOG* | Hs00750922_s1 | ThermoFisher | |
| *RNA45S5* | Hs03928990_g1 | ThermoFisher | |
| ***Site Directed Mutagenesis*** | | | |
| **Target Gene** | **Mutation** | **Forward** | **Reverse** |
| *ARHGEF26* | ∆PH domain (∆aa656-726) | CTCTTCCCGGTGGGGGAAGCCGCCTG | CAGGCGGCTTCCCCCACCGGGAAGAG |
| ***Cut and Paste Cloning*** | | | |
| **Target Gene** | **Source/Destination** | **Forward** | **Reverse** |
| *ARHGEF26* Constructs | pCMV Myc Constructs --> pEGFP3.1 | ATCGATCGATGTCGACGACTACAAGGACGACGATGACAAGATGGACGGCGAGAGCGAGGT | TTAAGCGCTATAGGATCCCTACACGTTGGTCTCCAGTC |
| ***Lambda-Red Recombination*** | | | |
| **Target Serovar** | **Target Gene** | **Forward for Cassette Generation** | **Reverse for Cassette Generation** |
| *S.* Typhimurium and *S.* Typhi | *sopB* | GAATGTTCCCACTCCCCTATTCAGGAATATTAAAAACGCTGTGTAGGCTGGAGCTGCTTC | ACGATTTAATAGACTTTCCATATAGTTACCTCAAGACTCACATATGAATATCCTCCTTAG |

| | | | |
|---|---|---|---|
| *S.* Typhimurium | *prgH* | CTGCTGCTATCGAGAACGACAGACATCGCTAACAGTATATGTGTAGGCTGGAGCTGCTTC | AAGGTGTTGCCATAATGACTTCCTTATTTACGTTAAATTACATATGAATATCCTCCTTAG |
| *S.* Typhi | *sopE* | ATATATAAATGAGTTATGTACATATAAAAGGATCATTACCGTGTAGGCTGGAGCTGCTTC | AGGAAGAGGCTCCGCATATTTTTTGGTTTTTCTGTGTTCACATATGAATATCCTCCTTAG |
| *S.* Typhi | *prgH* | CTGCTGCTATCGAGAACGACAGATATCGCTAACAGTATATGTGTAGGCTGGAGCTGCTTC | AAGATGTTGGCATAATGACTTCCTTATTTGCGTTAAATTACATATGAATATCCTCCTTAG |
| ***Strain Confirmation*** | | | |
| **Target Serovar** | **Target Gene** | **Forward** | **Reverse** |
| *S.* Typhimurium and *S.* Typhi | *sopB* | CCTGGTGCATAAAAGTCACATCC | CGGATTCATTAAAATAAACCTGTA |
| *S.* Typhimurium *and S.* Typhi | *prgH* | AATCCCTGTGTCCTGTGCGG | TATCCAGCGCCTCTGTTACC |
| *S.* Typhi | *sopE* | CATCAATCAGATGGACATAGCATTTGC | GACGGTTTAGCTCCGGAGTTAG |

1. Criss AK, Casanova JE. Coordinate regulation of Salmonella enterica serovar Typhimurium invasion of epithelial cells by the Arp2/3 complex and Rho GTPases. Infect Immun. 2003;71(5):2885-91.
2. Unsworth KE, Way M, McNiven M, Machesky L, Holden DW. Analysis of the mechanisms of Salmonella-induced actin assembly during invasion of host cells and intracellular replication. Cell Microbiol. 2004;6(11):1041-55.
3. Humphreys D, Davidson A, Hume PJ, Koronakis V. Salmonella virulence effector SopE and Host GEF ARNO cooperate to recruit and activate WAVE to trigger bacterial invasion. Cell Host Microbe. 2012;11(2):129-39.
4. Lilic M, Galkin VE, Orlova A, VanLoock MS, Egelman EH, Stebbins CE. Salmonella SipA polymerizes actin by stapling filaments with nonglobular protein arms. Science. 2003;301(5641):1918-21.
5. Hayward RD, Koronakis V. Direct nucleation and bundling of actin by the SipC protein of invasive Salmonella. EMBO J. 1999;18(18):4926-34.
6. Zhou D, Mooseker MS, Galan JE. An invasion-associated Salmonella protein modulates the actin-bundling activity of plastin. Proc Natl Acad Sci U S A. 1999;96(18):10176-81.
7. Francis CL, Starnbach MN, Falkow S. Morphological and cytoskeletal changes in epithelial cells occur immediately upon interaction with Salmonella typhimurium grown under low-oxygen conditions. Mol Microbiol. 1992;6(21):3077-87.
8. Stender S, Friebel A, Linder S, Rohde M, Mirold S, Hardt WD. Identification of SopE2 from Salmonella typhimurium, a conserved guanine nucleotide exchange factor for Cdc42 of the host cell. Mol Microbiol. 2000;36(6):1206-21.
9. Humphreys D, Davidson AC, Hume PJ, Makin LE, Koronakis V. Arf6 coordinates actin assembly through the WAVE complex, a mechanism usurped by Salmonella to invade host cells. Proc Natl Acad Sci U S A. 2013;110(42):16880-5.

10. Patel JC, Galan JE. Differential activation and function of Rho GTPases during Salmonella-host cell interactions. J Cell Biol. 2006;175(3):453-63.
11. Chen LM, Hobbie S, Galan JE. Requirement of CDC42 for Salmonella-induced cytoskeletal and nuclear responses. Science. 1996;274(5295):2115-8.
12. Ablain J, Xu M, Rothschild H, Jordan RC, Mito JK, Daniels BH, et al. Human tumor genomics and zebrafish modeling identify SPRED1 loss as a driver of mucosal melanoma. Science. 2018;362(6418):1055-60.
13. Patel JC, Galan JE. Investigating the function of Rho family GTPases during Salmonella/host cell interactions. Methods Enzymol. 2008;439:145-58.
14. Pujol C, Bliska JB. The ability to replicate in macrophages is conserved between Yersinia pestis and Yersinia pseudotuberculosis. Infect Immun. 2003;71(10):5892-9.
15. Wang RF, Kushner SR. Construction of versatile low-copy-number vectors for cloning, sequencing and gene expression in Escherichia coli. Gene. 1991;100:195-9.
16. Datsenko KA, Wanner BL. One-step inactivation of chromosomal genes in Escherichia coli K-12 using PCR products. Proc Natl Acad Sci U S A. 2000;97(12):6640-5.
17. Ellerbroek SM, Wennerberg K, Arthur WT, Dunty JM, Bowman DR, DeMali KA, et al. SGEF, a RhoG guanine nucleotide exchange factor that stimulates macropinocytosis. Mol Biol Cell. 2004;15(7):3309-19.
18. Awadia S, Huq F, Arnold TR, Goicoechea SM, Sun YJ, Hou T, et al. SGEF forms a complex with Scribble and Dlg1 and regulates epithelial junctions and contractility. J Cell Biol. 2019;218(8):2699-725.
19. He TC, Chan TA, Vogelstein B, Kinzler KW. PPARdelta is an APC-regulated target of nonsteroidal anti-inflammatory drugs. Cell. 1999;99(3):335-45.
